# *Nckx30c*, a *Drosophila* K^+^-dependent Na^+^/Ca^2+^ exchanger, regulates temperature-sensitive convulsions and age-related neurodegeneration

**DOI:** 10.1101/2025.10.08.681163

**Authors:** Al Nahian Khan Majlish, Sophia Bourgeois, Shu Hui Lye, Saul Landaverde, Emilia Cytron, Megan Cline, Harris Bolus, Robert N. Correll, Atulya Iyengar, Stanislava Chtarbanova

## Abstract

Calcium (Ca^2+^) homeostasis is fundamental to neuronal physiology, including in the regulation of membrane excitability and synaptic transmission. Disruptions in the ion transporters regulating Ca^2+^ influx and efflux are clearly linked to seizure disorders and age-related neurodegenerative disease. Yet, the specific contributions of variants in genes encoding these transporters to neurological disease remain to be fully understood. *Drosophila melanogaster* has proven to be a powerful genetic model for uncovering such mechanisms, particularly through studies of mutants that display temperature-sensitive (TS) behavioral phenotypes.

In a forward genetic screen, we identified a mutant line that exhibited TS convulsions along with progressive, age-dependent neurodegeneration. We mapped the mutation to *Nckx30c*, specifically within the transmembrane ion-binding region of this K^+^-dependent Na^+^/Ca^2+^ exchanger. Characterization of this mutant, together with a second *Nckx30c* allele, revealed TS convulsions, impaired locomotion, a markedly shortened lifespan, neurodegeneration with age, along with structural defects at larval and adult neuromuscular junctions (NMJs). *Nckx30c* mutants also displayed altered neural motor circuit performance. Gene expression analysis confirmed that *Nckx30c* levels were reduced in heads of *Nckx30c* loss-of-function mutants.

Tissue-specific manipulation revealed that knockdown of *Nckx30c* in neurons recapitulated the TS convulsions, locomotor defects, and shortened lifespan phenotypes.

*Drosophila Nckx30c* is highly conserved and shares homology with mammalian *SLC24A2*, a solute carrier family 24 member whose neurological role is not yet fully elucidated. Our work establishes *Nckx30c* as an essential regulator of neuronal health and provides an *in vivo* framework for investigating the contribution of *SLC24A2* to neuronal Ca^2+^ homeostasis, seizures and age-related neurodegeneration.

## Introduction

Neurodegenerative diseases (NDs) are a group of disorders characterized by the progressive loss of structure, function, and survival of neurons in the central nervous system. These conditions often lead to cognitive, motor, and behavioral impairments and are marked by common pathological features such as synaptic dysfunction and neuronal death. In 2019, it was estimated that NDs affect more than 57 million people worldwide, representing a major global health challenge. Most common examples include Alzheimer’s disease (AD) and Parkinson’s disease (PD) (Gadhave et al., 2024; Gao & Hong, 2008; Nichols et al., 2022; Peña-Bautista et al., 2020). One of the key features of NDs is neuronal dysfunction and cell death, which are often linked to disruption in calcium (Ca^2+^) homeostasis, though the underlying mechanism behind this link is still being actively investigated (Sun et al., 2024). Ca^2+^ functions as a critical second messenger in neurons, translating external stimuli into precise intracellular signals that regulate a wide array of physiological processes (Bagur & Hajnóczky, 2017; Berridge, 1998; Horigane et al., 2019; Pchitskaya et al., 2018). Ca^2+^ homeostasis is crucial for regulating various functions in the nervous system, including neuronal growth and differentiation, action potential generation, and the formation of learning and memory (M. Wang et al., 2025).

Disruption of Ca^2+^ dynamics not only compromises these essential processes but also contributes to neuronal hyperexcitability, which plays a central role in seizure generation (Xie et al., 2024). Seizures are episodes of sudden, abnormal, and hyper-synchronous neuronal firing that can result in transient disturbances in movement, behavior, sensation, or consciousness (Stafstrom & Carmant, 2015). This hyperexcitability is a hallmark of epileptic seizures and is often observed in neurodegenerative contexts (Amatniek et al., 2006; Gourmaud et al., 2022; Negi et al., 2023; Noebels, 2011; Tombini et al., 2021). When Ca^2+^ homeostasis is disrupted, abnormally elevated intracellular Ca^2+^ can lower the threshold for action potential initiation and enhance repetitive firing, resulting in neuronal hyperexcitability (Mahapatra et al., 2024; Zhou et al., 2022). Thus, altered Ca^2+^ signaling serves as a shared pathological mechanism bridging seizure and ND, emphasizing the importance of studying Ca^2+^ regulation in the context of both seizure disorders and NDs. Seizures are increasingly recognized as comorbid features in several NDs, including AD. AD patients, especially those with early-onset or familial forms, exhibit an elevated risk of both clinical and subclinical seizures (Lehmann et al., 2021; Purushotham et al., 2022).

In neurons, Ca^2+^ homeostasis is regulated by a coordinated system of different ion channels, transporters and pumps (Bkaily & Jacques, 2023; Brini et al., 2014; Britzolaki et al., 2018; Gleichmann & Mattson, 2011; Mozolewski et al., 2021; Nikoletopoulou & Tavernarakis, 2012; Pizzo et al., 2012). Under resting conditions, neurons maintain a steep Ca^2+^ gradient, with intracellular Ca^2+^ concentrations being approximately 10,000 fold lower than extracellular levels (Gleichmann & Mattson, 2011). This gradient is essential for rapid Ca^2+^ signaling upon stimulation, where the influx is maintained by different voltage gated Ca^2+^ channels during depolarization (Dolphin, 2021). To restore basal levels, excess Ca^2+^ is actively extruded by plasma membrane Ca^2+^-ATPases (PMCAs) (Krebs, 2022) and K dependent Na /Ca^2+^ exchangers (NCKXs) (A. H. Jalloul et al., 2020), the latter being a member of the most abundant Ca^2+^ extrusion protein family (Altimimi & Schnetkamp, 2007). NCKX transporters are receiving growing interest in the scientific community due to their emerging role as crucial regulators of Ca^2+^ signaling and homeostasis (Al-Khannaq & Lytton, 2022). These transporters are encoded by the *SLC24* gene family, which consists of five isoforms of the protein (NCKX1-5) found in mammals (Schnetkamp, 2013). They function with a distinct stoichiometry of exchanging 1 Ca^2+^ and 1 K out of the cell for 4 Na in, which allows for highly efficient Ca^2+^ extrusion, particularly under physiological conditions where Na and K gradients are maintained (Ali H. Jalloul et al., 2018). Among the five members of the group, NCKX2 is highly expressed in neurons, where it contributes to regulating calcium homeostasis and serves a protective role against Ca^2+^-induced excitotoxicity (Cuomo et al., 2022; Zhang et al., 2015).

Given the evolutionary conservation of calcium signaling mechanisms, model organisms like *Drosophila melanogaster* have proven invaluable for dissecting the roles of Ca^2+^ transporters in neural function (Chorna & Hasan, 2012). The genetic tractability, well-characterized nervous system, and availability of powerful tools for cell-type-specific gene manipulation make *Drosophila* an ideal system for studying ion transporters *in vivo* (Y. Wang et al., 2018).

In *Drosophila,* two genes encode NCKX: *zydeco* and *Nckx30c*. *zydeco*, is a well-characterized example supporting the importance of Ca^2+^ extrusion in brain function that encodes a Na /Ca^2+^, K^+^ exchanger expressed in glial cells (Winkfein et al., 2004). Disruption of *zydeco*’s function leads to impaired glial Ca^2+^ oscillations and seizures in *Drosophila* (Melom & Littleton, 2013). The *Nckx30c* was originally cloned and characterized in photoreceptor cells (Haug-Collet et al., 1999) but may have a broader role in regulating neuronal excitability and protecting against calcium-induced cytotoxicity in the brain, an area that remains underexplored. *Nckx30c* gene encodes a protein that has been described as a functional homolog of the NCKX family and is predicted to act as a K^+^-dependent Na /Ca^2+^ exchanger involved in Ca^2+^ homeostasis with a 71% sequence identity with mammalian NCKX2 (Haug-Collet et al., 1999; Webel et al., 2002).

Using a forward genetic approach and screening through an N-ethyl-N-nitrosourea (ENU)-mutagenized collection of *Drosophila* mutants for the second chromosome, we identified a novel mutant allele of *Nckx30c* that exhibits reversible TS convulsions with a point mutation that changes a threonine amino acid into a proline in the transmembrane domain of the protein. Notably, TS paralytic mutants are frequently associated with neurodegenerative phenotypes (Palladino et al., 2002), a pattern we also observed in aging *Nckx30c* mutants. In addition to these phenotypes, *Nckx30c* mutants exhibit impaired locomotion that progressively worsens with age. Tissue specific knockdown experiments using RNA interference (RNAi) showed that *Nckx30c* function is important in neurons unlike *zydeco,* which is glial specific (Melom & Littleton, 2013). In *Nckx30c* mutants, we also observed a morphological defect in the larval neuromuscular junction (NMJ) and loss of motor neuron branches in adult dorsal longitudinal muscle (DLM), suggesting abnormal synaptic function and neurotransmission. Based on its proposed primary function as a K^+^-dependent Na /Ca^2+^ exchanger, *Nckx30c* may be involved in maintenance of Ca^2+^ homeostasis in the cell, and disruption of this function may result in neuronal hyperexcitability and neurodegeneration. Further investigation into *Nckx30c* and related Ca^2+^ clearance mechanisms may provide valuable insights into the molecular pathways linking Ca^2+^ dysregulation to neurodegenerative disorders and seizure susceptibility.

## Materials and Methods

### *Drosophila* stocks and maintenance

*Drosophila* strains were maintained at 25°C on standard cornmeal medium. The *w^1118^* flies were a gift from Dr. John Yoder, University of Alabama. The ENU-mutagenized collection of *Drosophila* used in the screen including line 426 was a kind gift from Dr. Steven Robinow (University of Hawaii). This collection was previously used to screen for novel mutants that exhibit neurodegeneration (Loewen et al., 2018). The *w^1118^;Nckx30c^426^* stock was generated by backcrossing the 426 mutant line to *w^1118^* flies for at least 6 generations. Unless otherwise specified, young and aged cohorts were 4–7 days-old (labeled as 5-day-old) and 27–32 days-old (labeled as 30-day-old), respectively. The following fly lines were obtained from the Bloomington *Drosophila* Stock Center at Indiana University: *y^1^w^67c23^* (#6599), *w^1118^;Nckx30c^MB06102^* (#25318)*, w^1118^;Nckx30c^MB07279^*(#25268), deficiency lines for *Nckx30c* – *Df(2L)s1402* (#556), *Df(2L)Exel6024* (#7507), *Df(2L)ED695* (#8041), and for RNAi knockdown experiments, *elav-Gal4;UAS-dcr2* (#25750)*, repo-Gal4* (#7415), four *UAS-Nckx30cRNAi* lines (#27246), (#34385), (#34386) and (#42581). An institutional ethics approval was not required for this study.

### Lifespan analysis

Flies were collected three days post-eclosion and maintained at 25°C. Cohorts were transferred to fresh vials every two to three days, and daily after 20-30 days of age, to prevent older flies from adhering to the food. Each vial contained 25 flies of the same genotype. The number of surviving flies was recorded daily. Three independent survival experiments were performed for each genotype under identical conditions.

### Temperature sensitive paralysis assay

For TS paralysis assay flies were collected at three days post-eclosion and aged at 25°C until they were 4-7 days old. Flies were separated by sex during the collection with 20-25 flies in each vial. The female flies were transferred to fresh vials if the food became moist or larvae were observed. For the assay flies were transferred to glass vials and placed into a glass water tank preheated to 38°C and the number of flies paralyzed or exhibiting convulsive behavior in 2 minutes was recorded. Convulsive behavior refers to the uncontrolled jerking of legs and the inability to move or fly properly (Dare et al., 2020). Paralytic behavior is defined as the fly lying on its back with little to no movement of legs and wings (Williamson et al., 1974). For the TS paralysis assay done as part of the forward genetic screen, flies were raised and aged at 25°C but transferred and kept at 18°C for 24 hours prior to TS testing.

### Automated locomotion assay (video tracking)

Fly locomotion was measured using an automated open-field tracking system with heating arena following the protocol described in (Landaverde et al., 2024). Flies were placed in polyacrylate arenas set on Whatman #1 filter paper, which rested on a Peltier stage (AHP-1200CPV, TECA Corporation). Temperature within the arena was tracked using a T-type thermocouple (5SRTC-GG-T-30-36, Omega Engineering) connected to a National Instruments NI TC-01 data acquisition card, while stage temperature was regulated by a custom LabVIEW program. Lighting was supplied by a PVC cylinder fitted with LED strips and intensity was kept at ∼1000 lux with a current controller (∼100 mA). Groups of four flies of the same genotype and sex were loaded into the arenas without anesthesia and recorded using a Logitech C920 webcam (30 fps). The assay protocol included baseline at 22°C (180 s), high-temperature at 36-39°C (180 s), and recovery at baseline temperature (180 s). Locomotion recordings were controlled by a LabVIEW script and analyzed in MATLAB (R2024b, Mathworks,) with IowaFLI Tracker to calculate distance traveled (Iyengar et al., 2012).

### Immunohistochemistry

For larval NMJ analysis, wandering third instar larvae were dissected in phosphate buffered saline (1X PBS). After removing internal organs, specimens were fixed in 4% paraformaldehyde (PFA) in 1X PBS for 30 minutes on a rotating shaker. After removal of fixative solution, specimens were washed 2 times with 0.1% Triton X-100 in 1X PBS (PBS-T) for 10 minutes each, replacing with fresh PBS-T at each step. After the final wash larvae were incubated in a blocking solution of PBS-T and 4% normal goat serum (MP Biomedicals, Cat. # ICN19135680) for 1 hour. After removing the blocking solution, the larvae were incubated for another hour with 1:200 Alexa Fluor 594 goat anti-horseradish peroxidase (Jackson ImmunoResearch Cat. #123-585-021). Incubation with the antibodies was done in dark conditions and at room temperature.

After the incubation, the antibodies were removed, and the larvae were washed 3 times with PBS-T for 10 minutes each on a rotating shaker. Following the final wash, samples were mounted in ProLong™ Glass Antifade Mountant with NucBlue™ (Invitrogen, Cat. # P36981). Imaging was done using Nikon Eclipse Ti2 Laser Scanning Confocal Microscope and processed using Fiji ImageJ2 (Version: 2.16.0/1.54p). Boutons and satellite boutons were separately counted.

For dorsal longitudinal muscle (DLM) NMJ analysis, DLMs were prepared by removing the head, wings, legs and abdomen from adult thoraces, followed by fixation in 4% paraformaldehyde in 1X PBS for 30 minutes at room temperature. Thoraces were washed 4 times with 1X PBS and flash-frozen by placing the tubes with thoraces in liquid nitrogen for 20 seconds. Frozen thoraces were bisected along the midline in ice-cold 1X PBS and incubated in blocking buffer of PBS-T and 4% normal goat serum (MP Biomedicals, Cat. # ICN19135680) for at least 1 hour at 4°C, adapted from the protocol of Sidisky and Babcock (2020) (Sidisky & Babcock, 2020). Bisected hemi-thoraces were incubated with 1:200 Alexa Fluor 594 goat anti-horseradish peroxidase (Jackson ImmunoResearch, Cat. # 123-585-021) in blocking solution for 2 hours in the dark at room temperature. Following staining, samples were washed 4 times in PBS-T for 5 minutes each. Samples were mounted in Invitrogen™ ProLong™ Glass Antifade Mountant with NucBlue™ (Invitrogen, Cat. # P36981). Imaging was done using Nikon Eclipse Ti2 Laser Scanning Confocal Microscope and processed using Fiji ImageJ2 (Version: 2.16.0/1.54p).

### Quantification of neuronal branching

Neuronal branching complexity was quantified from confocal images using the Analyze Skeleton (2D/3D) plugin in Fiji ImageJ2 (Version: 2.16.0/1.54p) (Arganda-Carreras et al., 2010). Only high-quality images with well-preserved muscle fibers and clearly resolved neurons were included in the analysis. Maximum intensity projection images were first converted to 8-bit grayscale, and the signal threshold was adjusted to isolate neuronal arbors from the background. Then they were converted to binary format and skeletonized to generate a one-pixel-wide representation of the neuronal structure. Skeletonized images were analyzed using the plugin function available under the Analyze Skeleton menu in Fiji. The “Prune cycle method” was set to “None” to ensure accurate branch tracing without removal of loops. The plugin automatically quantified the total number of branches, junctions, end-point voxels, and the average branch length for each neuron (Young & Morrison, 2018). For each image, the area of measurement was 45000 μm^2^. To analyze the branch point ratio, the number of junctions was divided by the total of junction numbers and endpoint numbers. This reveals the branching complexity of the analyzed image. These parameters provided quantitative measures of neuronal arbor complexity and were used to compare genotypes statistically (Scholtens et al., 2022).

### Histology

Flies were collected upon eclosion and aged at 29°C. Severed fly heads were placed in fresh Carnoy’s fixative solution (6 parts 100% ethanol, 3 parts chloroform, and 1 part glacial acetic acid) and left overnight at 4°C before exchanging into 70% ethanol the next day. Then the heads were processed into paraffin blocks using standard histological procedures following the protocol as described in Gevedon *et al*. (Gevedon et al., 2019). Head samples were paraffin embedded at the Experimental Animal Pathology Lab (EAPL) facility at the University of Wisconsin-Madison. Using a microtome (Leica Biosystems), embedded heads were sectioned at 5 μm and stained using hematoxylin and eosin (H&E). Neurodegeneration was observed on the stained midbrain sections using a light microscope (Nikon) and quantified based on neurodegeneration index as described by Cao *et al*. (Cao et al., 2013). The average of three consecutive midbrain slices were used to assign the final score of one brain sample.

### Tethered fly electrophysiology

The procedure for tethered fly electrophysiology was adapted from Iyengar & Wu (2014) and Landaverde et al. (2024) (Iyengar & Wu, 2014; Landaverde et al., 2024). 5-day-old flies were anesthetized on ice and secured onto a tungsten pin using UV-cured cyanoacrylate glue. After a 10-minute recovery period, electrolytically sharpened tungsten electrodes were placed into the top-most dorsal longitudinal muscle (DLMa) and into the dorsal abdomen (for reference). Muscle action potentials were amplified via an AC amplifier (AM Systems #1800) and digitized by a data acquisition card (National Instruments USB 6210) controlled by a custom-written LabVIEW script. Signals were analyzed off-line using custom-written MATLAB scripts (Landaverde et al., 2024).

Stimulation of the giant-fiber jump and flight escape circuit was performed through sharpened tungsten electrodes inserted into the left and right cornea driven by an AM systems (#2100) stimulator controlled by the data acquisition card.

To gradually heat flies, air was pumped (flow rate = 4 l/min, Tetra Whisper 60 pump) through a bubble humidifier containing deionized water and passed through a process heater (Omega AHP 3741). The outlet of the heater (diameter: ∼ 1 cm) was placed ∼ 3 cm from the tethered fly. Air temperature exiting the heater was measured by a T-type thermocouple and continuously digitized by a TC-01 temperature input device (National Instruments). The heater was driven by a current controlled DC power supply. During the protocol, the heater generated 11.88 W of heat for the first 180 s, followed by 11 W of heat for the next 120 s. After the 300-s protocol, both the air pump and heaters were shut off.

### Gene mutation analysis

Genomic DNA was extracted from lines *1484* (a line from the ENU mutagenized collection not exhibiting TS paralysis at 4-7 days of age), *426*, and *y^1^w^67c23^* whole flies using the PureLink Genomic DNA Mini Kit (Invitrogen) following manufacturer’s instructions. Primers targeting *Nckx30c* exons (**Supplemental Table 1**) were used for amplification by PCR with Phusion® High-Fidelity DNA Polymerase (Thermo Scientific™) and DMSO, following the manufacturer’s recommendations. PCR cycling conditions were as follows: initial denaturation at 98°C for 30 seconds; 35 cycles of denaturation at 98°C for 10 seconds, annealing at 65°C for 30 seconds, and extension at 72°C for 35 seconds; followed by a final extension at 72°C for 1 minute and 15 seconds. PCR product sizes were confirmed by electrophoresis on a 1% agarose gel using the GeneRuler 1 kb Plus DNA ladder (Thermo Fisher).

For cDNA analysis, total RNA was extracted from whole flies of the *426* and *y^1^w^67c23^*strains. cDNA synthesis was performed using the SuperScript™ IV First-Strand cDNA Synthesis Kit (Invitrogen) with a *Nckx30c*-specific primer (**Supplemental Table 1**), according to the manufacturer’s instructions. PCR amplification of cDNA was carried out under the following conditions: initial denaturation at 98°C for 30 seconds; 35 cycles of denaturation at 98°C for 10 seconds and annealing/extension at 75°C for 2 minutes and 15 seconds; followed by a final extension at 75°C for 10 minutes. Sanger sequencing for both genomic and cDNA PCR products was performed by Eurofins Genomics, and sequence data were analyzed using Unipro UGENE software (Okonechnikov et al., 2012).

For predicting secondary protein structure organization, we used PSIPRED (Protein Analysis Workbench) (McGuffin et al., 2000). Using the full-length *Drosophila Nckx30c* amino acid sequence, we ran the web server with default settings, which returns per-residue probabilities for helix (H), strand (E), and coil (C).

### Gene expression analysis

Total RNA was extracted from the heads of 15-20 flies per genotype (7 day-old) using the Quick-RNA MicroPrep Kit (Zymo Research), following the manufacturer’s instructions. cDNA was synthesized from 500 ng of the extracted RNA using the High-Capacity cDNA Reverse Transcription Kit (Applied Biosystems).

RT-qPCR reactions were performed using SYBR Green Master Mix (Applied Biosystems) in a 10 µL total volume. Each reaction contained 5 µL SYBR Green, 0.5 µL each of 10 µM forward and reverse primers, and nuclease-free water to make up 9 µL of reaction mix. 1 microliter of cDNA, diluted 1:10 with nuclease-free water, was added to each reaction well.

Amplification was carried out using a standard two-hour RT-qPCR protocol. The primers used for RT-qPCR reactions are listed in the **Supplemental Table 1.**

## Statistical analysis

All statistical analyses were performed in GraphPad Prism v.10, except for automated locomotion data, where statistical analysis was done in MATLAB using the statistics toolkit. For each test, the “n” numbers are given in the figure legends. Lifespan (survival) curves were compared using the Log-Rank (Mantel-Cox) test. TS convulsions data were analyzed using one-way ANOVA with Tukey’s multiple comparisons. Automated locomotion data were analyzed with a Kruskal-Wallis test followed by a Bonferroni-corrected rank-sum post hoc test. For larval NMJ bouton and satellite bouton counts, mean values were assessed by one-way ANOVA with Dunnett’s post-hoc test versus the control. Histology based neurodegeneration quantification was analyzed by two-way ANOVA with Tukey’s multiple comparisons. For gene expression analysis, relative expression was calculated by the 2^-ΔΔ*CT*^ method using *Rp49* as the endogenous control and relative to *w^1118^* (set to 1), and group differences were tested by one-way ANOVA with Dunnett’s post-hoc test. For *Nckx30c* knockdown confirmation analysis, expression data were normalized to the *Rp49* gene as an endogenous control, and each knockdown was also normalized to its cell-type control. *elav-Gal4;UAS-dcr2>UAS-Nckx30cRNAi* was normalized to *elav-Gal4;UAS-dcr2>*+ and *repo-Gal4>UAS-Nckx30cRNAi* was normalized to *repo-Gal4>*+.

## Results

### Identification and mapping of a novel mutant exhibiting TS convulsions

To identify novel genes involved in temperature-sensitive (TS) convulsions and neurodegeneration in *Drosophila*, we performed a TS paralysis screen using 50 lines from a larger collection of ENU mutant *Drosophila* for chromosome 2. Among the 50 mutant lines tested, the line designated as *426* exhibited 100% paralysis or convulsions at 38°C in 5-day-old flies (**Figure 1A**). The convulsive phenotype was only observed in homozygous mutants, whereas heterozygous *426* flies (obtained by crossing *426* mutants with *Canton S wild type* flies), displayed wild type phenotype, suggesting a recessive nature of the mutation (**Figure 1B**). To map the location of the mutation on *Drosophila* chromosome 2, we performed complementation tests with a series of deficiency lines (8 deficiency lines were used and crossed with the *426* mutant) based on TS convulsive behavior. A schematic showing the cytological location for each deficiency line used is shown in **Supplemental Figure S1.** The mutant line *426* failed to complement the deficiency line *Df(2L)N22-14* in which cytological region 29C1-30C9 is deleted (**Figure 1C**). In addition to *Df(2L)N22-14*, we crossed five other deficiency lines spanning this region to *426* mutants. This allowed us to more precisely identify the region on chromosome 2L where the TS phenotype overlapped. Consequently, we mapped the TS phenotype to the left arm of chromosome 2 (30C6-30C7 cytological region), where the gene *Nckx30c* is located (**Supplemental Figure S2**). Henceforth, *426* is referred to as *Nckx30c^426^*.

**Figure 1:**
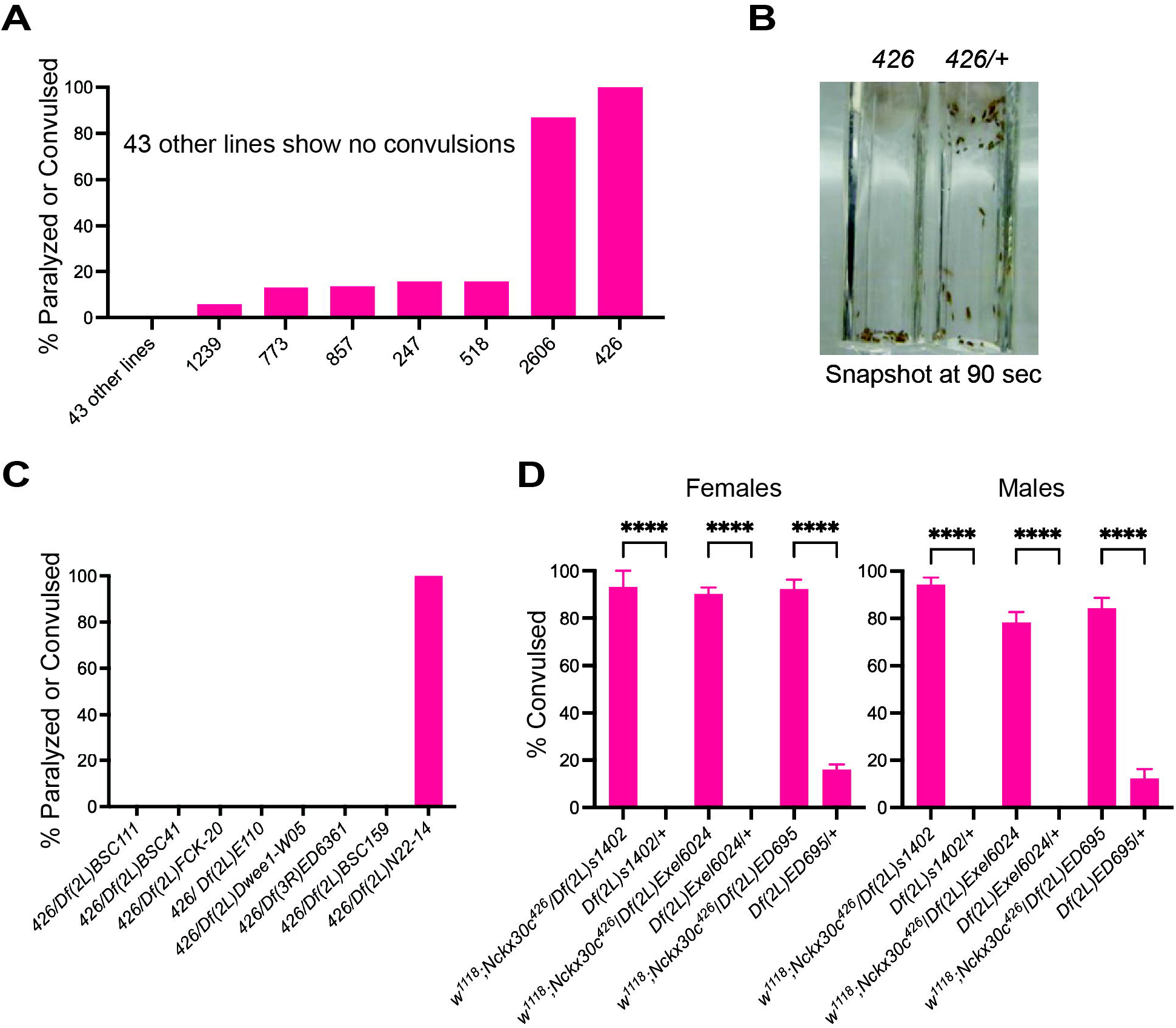
Phenotypic characterization of mutant 426 exhibiting temperature sensitive convulsions. **(A)** TS paralysis screen of 50 ENU-mutagenized chromosome 2 lines 5-day-old flies from each line were placed in a 38°C water bath, and the percentage of flies showing paralysis or convulsion within 5 min was calculated. Lines showing > 0% paralysis or convulsion are presented by the graph bars. **(B)** Representative image of the TS paralysis assay at 90 seconds shows convulsive behavior in homozygous *426* mutants compared to heterozygous control (*426* mutant crossed with *Canton S*). **(C)** Complementation tests using deficiency lines. Mutant *426* failed to complement *Df(2L)N22-14* and showed TS convulsions. Here, *Df(3R)ED6361* was used as a negative control. **(D)** Complementation tests for both females and males, using additional deficiency lines, refined the candidate region. Here, controls are heterozygotes obtained by crossing each deficiency line to *w^1118^*. Bars in the graph indicate the percentage of flies convulsed and mean ± SEM. For each line tested, the population size is n = 50. Statistics: one-way ANOVA with Tukey’s post hoc test; \*\*\*\**P* < 0.0001. All the specific P values are reported in the **Supplemental datasheet 1.**

Moreover, we performed complementation tests using three additional deficiency lines *Df(2L)s1402, Df(2L)Exel6024 and Df(2L)ED695* in this chromosomal region for both females and males. *Nckx30c^426^* mutant failed to complement all three deficiency lines in both sexes. Each deficiency line was compared with its control, which was crossed with *w^1118^*. This further confirmed the mapped candidate location (**Figure 1D).** The mapping for these deficiency lines is shown in **Supplemental Figure S3**.

### *Nckx30c^426^* mutants carry a point mutation in exon 2

We performed Sanger sequencing on the *Nckx30c^426^* mutant, using the *1484* line (from the TS screen and not showing the TS phenotype at the age at which flies were examined) and *y^1^w^67c23^* flies as controls. We sequenced the coding regions spanning all eight exons of *Nckx30c* and compared them with the reference DNA sequence. Our analysis revealed a single nucleotide substitution at position 13,179 in exon 2, where adenine (A) was replaced by cytosine (C) in the *Nckx30c^426^* mutant but not in the controls (**Figure 2A**). We also performed cDNA sequencing to confirm that the A to C substitution was retained in the mature mRNA and not removed through alternative splicing. Subsequent amino acid analysis showed that this mutation results in a missense change, replacing threonine (T) with proline (P) (**Supplemental Figure S4A**). By using PSIPRED prediction tool for predicting helix regions in protein structure, we found that the threonine lies in the alpha helix region of the protein (**Supplemental Figure S4B**). This amino acid substitution is predicted to disrupt helix structure. We used the Pfam (currently InterPro) database to analyze the mutant amino acid sequence and found that the T to P change was in the Na^+^/Ca^2+^ exchange domain (**Figure 2A**). Additionally, we analyzed the structure of the mutant protein using SWISS-MODEL, a web-based automated protein structure homology-modelling program (Bertoni et al., 2017; Bienert et al., 2017; Guex & Peitsch, 1997; Studer et al., 2020; Waterhouse et al., 2018) and compared it with the reference *Nckx30c* structure (**Figure 2B**). We found that the residue change of T to P was indeed predicted to be in the transmembrane domain. Multiple alignments of the Nckx30c protein sequence including the mutant amino acid against other *Drosophila* species and mammalian (*Homo sapiens* and *Mus musculus*) NCKX2 showed that this region is conserved among all the species (**Figure 2B**).

**Figure 2:**
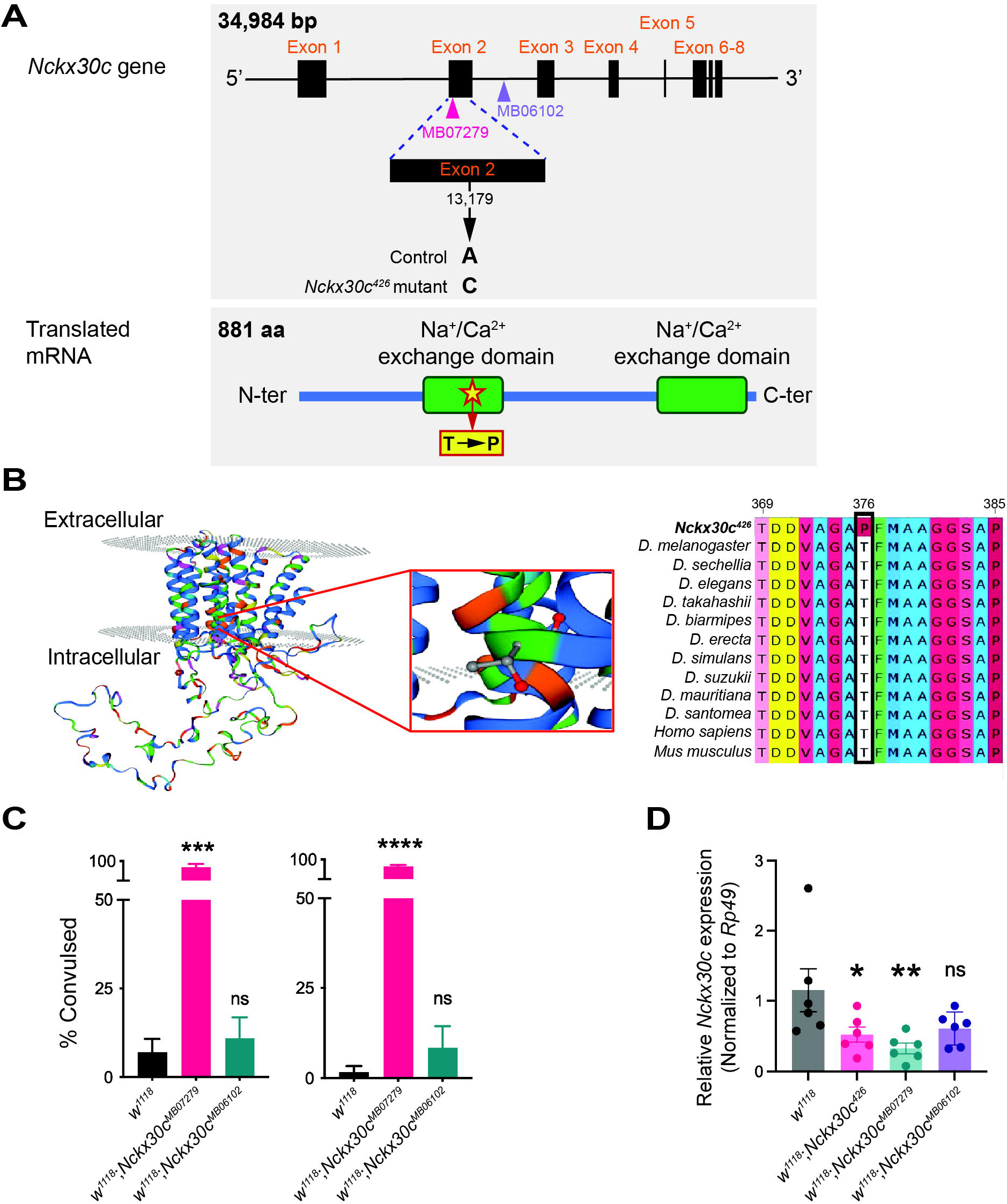
Molecular characterization of the *Nckx30c* mutant. **(A)** Schematic of the *Nckx30c* gene structure and associated mutations. The *426* mutant carries a point mutation (A to C) in exon 2, affecting a conserved region within the Na^+^/Ca^2+^ exchange domain. The location of two additional transposable element insertions (*MB07279* (exon 2)) and (*MB06102* (intron 2), are shown. The polypeptide corresponding to Nckx30C-PD is shown. **(B)** The predicted protein structure of *Nckx30c^426^* showing the mutation located in the transmembrane region. The inset (magnified region) highlights the location of the mutation. The affected region (residue 376) is evolutionarily conserved across different *Drosophila* species (10 species) and mammalian NCKX2 (*Homo sapiens* and *Mus musculus*), as shown by the amino acid sequence alignment of Nckx30c. Here, threonine (T) is replaced by proline (P) in the mutant. **(C)** TS paralysis assay of the P-element insertion alleles *w^1118^;Nckx30c^MB07279^* and *w^1118^;Nckx30c^MB06102^* of both females and males compared against control *w^1118^*. For each line tested, the population size are: Females: *w^1118^* n = 47, *w^1118^;Nckx30c^MB07279^* n = 43 and *w^1118^;Nckx30c^MB06102^* n = 50. Males: *w^1118^* n = 55, *w^1118^;Nckx30c^MB07279^* n = 53 and *w^1118^;Nckx30c^MB06102^* n = 55. The bar graphs show mean ± SEM; the symbols over each mutant genotype bar indicate statistical comparison against *w^1118^*. Statistics: one-way ANOVA with Dunnett’s post hoc test; \*\*\**P* ≤ 0.001, \*\*\*\**P* ≤ 0.0001, *ns* = not significant. All the specific P values are reported in the **Supplementa datasheet 1. (D)** RT-qPCR analysis of relative expression of *Nckx30c* gene in *w^1118^;Nckx30^426^*, *w^1118^;Nckx30c^MB07279^* and *w^1118^;Nckx30c^MB06102^*, compared to *w^1118^* controls. The data presented here were calculated using the 2^-ΔΔ*CT*^ method. Data were normalized to *Rp49* gene as an endogenous control and *w^1118^* as the control strain. The bar graphs show mean ± SEM; the symbols over each mutant genotype bar indicate statistical comparison against *w^1118^*. Statistics: one-way ANOVA with Dunnett’s post hoc test; \**P* ≤ 0.05, \*\**P* ≤ 0.01, *ns* = not significant. All the specific P values are reported in the **Supplemental datasheet 1.**

To determine whether disruption of *Nckx30c* triggers the TS and neurodegenerative phenotypes, and to evaluate how mutations in different parts of the gene impact its function, we utilized two additional transposable element insertion mutants: *w^1118^*;*Nckx30c^MB07279^*, (insertion in exon 2) and *w^1118^*;*Nckx30c^MB06102^* (insertion in intron 2) (**Figure 2A**). TS paralysis assays revealed that *w^1118^*;*Nckx30c^MB07279^*but not *w^1118^*;*Nckx30c^MB06102^* mutant exhibited a convulsive phenotype similar to the *Nckx30c^426^* mutant (**Figure 2C**). In the *w^1118^*;*Nckx30c^MB07279^* mutants, 92% of females and 88% of males exhibited the TS convulsion phenotype. In contrast, the *w^1118^*;*Nckx30c^MB06102^* mutants showed the phenotype in only 13% of females and 8% of males. Gene expression analysis from the heads of 7-day-old males of these mutant lines (**Figure 2D**) demonstrated that *w^1118^*;*Nckx30c^426^* displayed significantly reduced *Nckx30c* expression compared to the *w^1118^* control (*P* =0.0477). Additionally, *w^1118^*;*Nckx30c^MB07279^* displayed even more reduction in expression than *w^1118^* (*P* =0.0087). In contrast, *w^1118^*;*Nckx30c^MB06102^* didn’t show statistically significant differences relative to *w^1118^*. Together, the sequencing of the *Nckx30c^426^* and the characterization of the independent transposable element alleles suggest that the loss-of-function or reduced expression can produce the TS convulsion phenotype.

### *Nckx30c* mutants exhibit age-dependent neurodegeneration and shorter lifespan

Temperature-sensitive (TS) paralytic mutants are often associated with neurodegeneration (Palladino et al., 2002). In *Drosophila*, paralytic and convulsive behavior are both phenotypes of the nervous system abnormalities, sharing some common mechanisms (Burg & Wu, 2012). A secondary histology screen using paraffin-embedded brain sections of 4-week-old *Nckx30c^426^* mutants (n=3) and *y^1^w^67c23^* (n=3) controls was tested for manifestation of neuropathology (**Supplemental Figure S5**). We observed the appearance of holes, considered a hallmark of neurodegeneration, in mutant but not in control brains. Here, it needs to be noted that these images were obtained at the initial stages of characterization of the *Nckx30c^426^*mutant, and quantification was not done. Given that *w^1118^*;*Nckx30c^MB07279^*also exhibited TS convulsions, we further examined neurodegeneration in this mutant by comparing brain morphology between *w^1118^*;*Nckx30c^MB07279^*and control *w^1118^* flies at 5 and 30 days of age **(Figure 3A)**.

**Figure 3:**
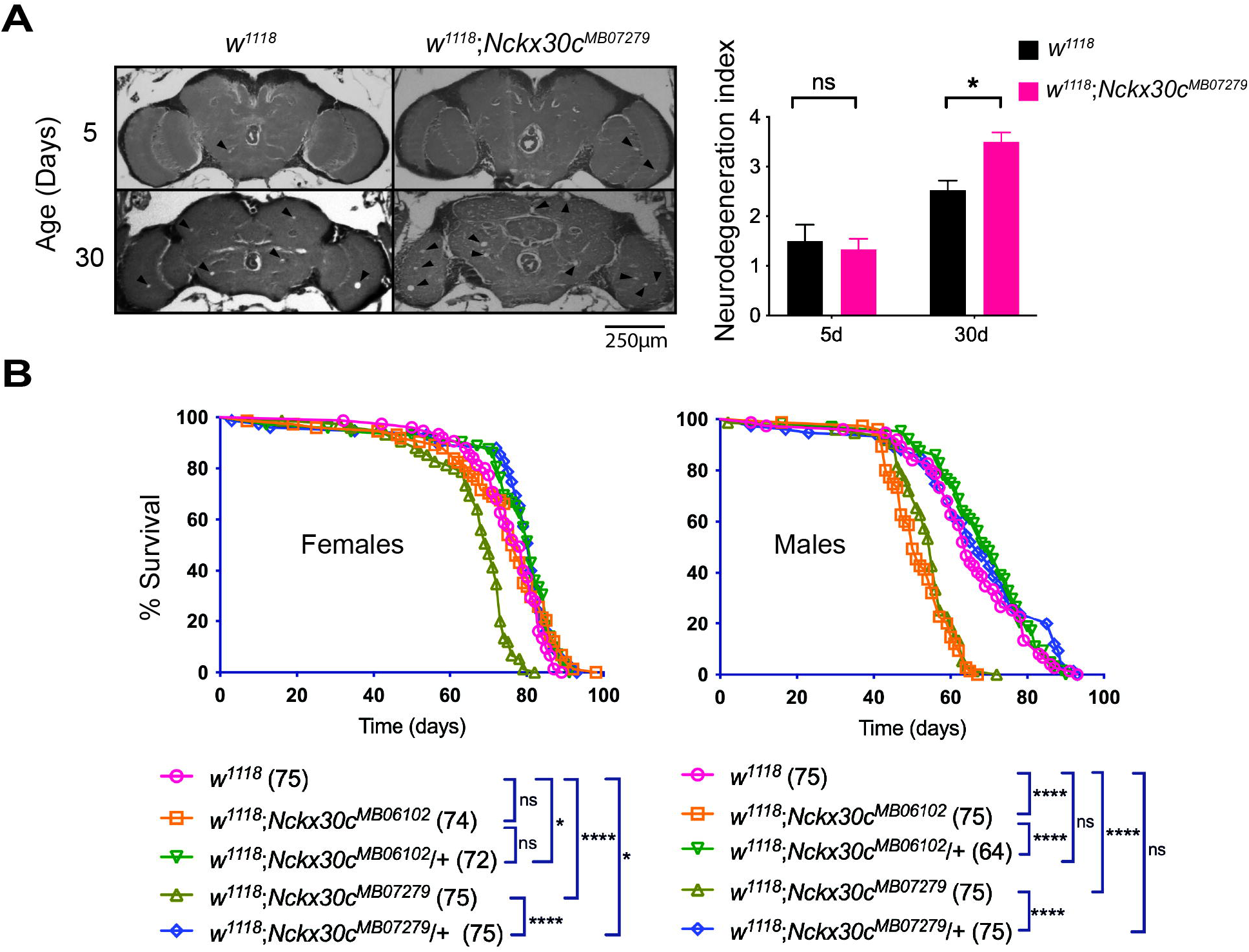
***Nckx30c* mutants exhibit age-dependent neurodegeneration and shorter lifespan. (A)** Left panel: representative 5-μm paraffin sections at approximately midbrain of *w^1118^* and *w^1118^;Nckx30c^MB07279^*. Arrowheads indicate holes in the brain indicative of neurodegeneration. Right panel: neurodegeneration index of young (5d) and old (30d) *w^1118^;Nckx30c^MB07279^* mutants and *w^1118^* controls. Scale bar represents 250 μm. Quantification of neurodegeneration in fly brains done based on the neurodegeneration index scale from 0-5 as in Cao *et al*. (2013). Values shown are mean ± SEM; two-way ANOVA with Tukey’s multiple comparison test; \**P* < 0.05, *ns* = non-significant. **(B)** Lifespan curves for flies of the indicated genotypes maintained at 25°C. Numbers in parentheses indicate sample size. Log-Rank (Mantel-Cox); \**P* ≤ 0.05, \*\*\*\**P* ≤ 0.0001, *ns* = not significant. All the specific P values are reported in the **Supplemental datasheet 1**.

Neurodegeneration was quantified in a double-blind manner using the neurodegeneration index described by Cao *et al*. (Cao et al., 2013) where higher scores indicate more severe degeneration (see Methods section for details). For 5 day-old flies, *w^1118^*;*Nckx30c^MB07279^* (n=6) and *w^1118^* (n=8) displayed comparable neurodegeneration indexes of 1.3 and 1.5 respectively, which were not significantly different (*P* =0.978). However, by 30 days, *w^1118^*;*Nckx30c^MB07279^*brains (n=8) exhibited a significantly higher score of 3.5 compared to *w^1118^*controls (n = 19) scoring 2.5 (*P* = 0.0243). This indicates that *w^1118^*;*Nckx30c^MB07279^* mutants show age-dependent neurodegeneration in comparison to controls.

We next assessed the lifespan of both homozygous *w^1118^*;*Nckx30c^MB07279^* and *w^1118^*;*Nckx30c^MB06102^*, as well as heterozygous *w^1118^*;*Nckx30c^MB07279^*/+ and *w^1118^*;*Nckx30c^MB06102^*/+ mutants and compared these flies to *w^1118^* controls **(Figure 3B)**. In females, *w^1118^*;*Nckx30c^MB07279^*(n=75) displayed significantly shorter lifespan compared to *w^1118^*;*Nckx30c^MB07279^*/+ (n=75); *P* <0.0001 and *w^1118^* (n=75); *P* <0.0001 where nearly all the flies died around 80 days. Female *w^1118^*;*Nckx30c^MB06102^* (n=74) didn’t show any significant change in their lifespan compared to *w^1118^*;*Nckx30c^MB06102^*/+ (n=72) and *w^1118^* flies. Both the heterozygotes *w^1118^*;*Nckx30c^MB07279^*/+ and *w^1118^*;*Nckx30c^MB06102^*/+ lived longer than control *w^1118^* (*P* =0.0191 and *P* =0.0289 respectively).

Similarly, in males, *w^1118^*;*Nckx30c^MB07279^*(n=75) displayed significantly shorter lifespan compared to *w^1118^*;*Nckx30c^MB07279^*/+ (n=75); *P* <0.0001 and *w^1118^* (n=75); *P* <0.0001 where almost all homozygous mutants died around 70 days. However, male *w^1118^*;*Nckx30c^MB06102^*(n=75) also showed significantly shorter lifespan than *w^1118^*;*Nckx30c^MB06102^*/+ (n=64); *P* <0.0001 and *w^1118^* (*P* <0.0001) flies. For males, the heterozygotes didn’t show any significant change in their lifespan compared to the *w^1118^* strain. These findings show that homozygous *w^1118^*;*Nckx30c^MB07279^* mutants exhibit a significantly reduced lifespan in both males and females, whereas only male homozygous *w^1118^*;*Nckx30c^MB06102^*mutants display a shorter lifespan. This indicates that some other internal processes in males reduce the lifespan, as we didn’t observe a significant reduction of *Nckx30c* expression in *w^1118^*;*Nckx30c^MB06102^*male fly heads. During the early stages of characterization of the *426* mutants, we also observed a significantly reduced lifespan in homozygous *Nckx30c^426^* compared against the laboratory control strain *y^1^w^67c23^* and heterozygous *Nckx30c^426^*/+ flies (**Supplemental Figure S6**). This was observed in both females and males.

### Locomotor deficits in *Nckx30c* mutants exacerbate with aging and at higher temperatures

To evaluate locomotor activity, we used an automated movement tracking system to record flies carrying different *Nckx30c* mutant alleles. Representative movement tracks at 22°C illustrate the locomotor defects in *Nckx30c* mutants compared with the *w^1118^*controls (**Figure 4A**). At 5 days, both *w^1118^*;*Nckx30c^426^*and *w^1118^*;*Nckx30c^MB07279^* mutants already showed reduced activity compared to *w^1118^* and *w^1118^*;*Nckx30c^MB06102^*flies, and these defects became more severe by 30 days of age.

**Figure 4:**
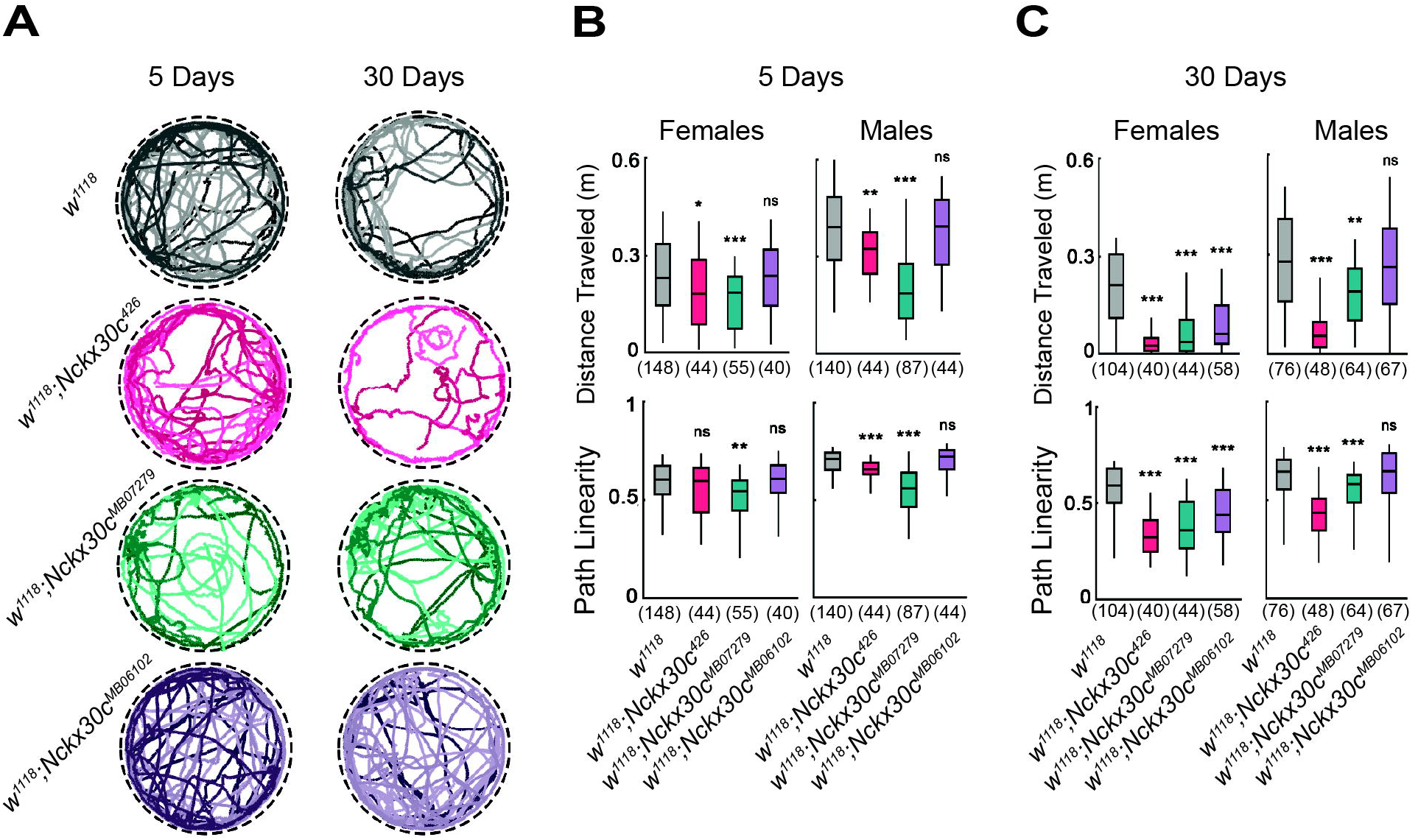
**Automated video tracking of locomotor activity in *Nckx30c* mutants at 22**°**C. (A)** Representative tracks of four male flies from the respective genotype (Temperature: 22°C). Left panel: 5-day-old flies, right panel: 30-day-old flies. **(B, C)** Distance traveled and path linearity at 22°C in females and males flies at 5 and 30 days were measured. Each set of box plots represents each genotype. Numbers in parentheses represent sample size. Statistics: Kruskal Wallis non-parametric ANOVA (Bonferroni-corrected rank-sum post hoc test). Data from each mutant genotype were compared against *w^1118^* to generate statistical significance, represented by the symbols over each box. \*\**P* ≤ 0.01, \*\*\**P* ≤ 0.001, *ns* = not significant. All the specific P values are reported in the **Supplemental datasheet 1**.

Quantification of total distance traveled over a 3-minute observation period at 22°C was measured in both young (5 days old) and aged (30 day old) flies, for both sexes (**Figure 4B**). At 5 days of age, female *w^1118^*;*Nckx30c^426^*flies on average traveled 0.18 m which 21% less than *w^1118^* females (0.23 m), while male *w^1118^*;*Nckx30c^426^*travelled 0.32 m which is 17% less in comparison to *w^1118^* males (0.39 m). Path linearity, defined as the ratio of displacement to total distance over 2-second intervals, was also reduced in both males and females (data available in **Supplemental datasheet 2**). In comparison to *w^1118^* controls, *w^1118^*;*Nckx30c^MB07279^*flies displayed a similar pattern, with reductions in distance traveled of 20% (females: 0.18 m) and 53% (males: 0.18 m), and reductions in path linearity of 10% (females) and 21% (males). By contrast, *w^1118^*;*Nckx30c^MB06102^*flies did not differ significantly from *w^1118^* at this age.

At 30 days of age, all genotypes exhibited lower locomotion than their younger counterparts (**Figure 4C**). In *w^1118^*;*Nckx30c^426^* flies, distance traveled was reduced by 88% in females (0.02 m) and 80% in males (0.05 m), compared to *w^1118^* (females: 0.20 m and males: 0.27 m). Path linearity declined by 45% and 33%, respectively, compared with *w^1118^*. *w^1118^*;*Nckx30c^MB07279^* flies also showed pronounced locomotor impairment, with distance traveled reduced by 83% (females: 0.03 m) and 32% (males: 0.18 m), and path linearity reduced by 39% (females) and 10% (males). For *w^1118^*;*Nckx30c^MB06102^*, we observed a significant reduction in distance traveled only in 30-day-old females (71%, 0.06 m), whereas males did not differ significantly from controls. Path linearity remained unchanged in both sexes for this genotype (data available in **Supplemental datasheet 2**).

We evaluated locomotor activity in *Nckx30c* mutants during a ramp to high temperature (39°C) to see how elevated temperature affects motor function. Representative movement tracks of a single male fly at 39°C (**Figure 5A**) and speed profiles of four flies (**Figure 5B)** contrast locomotor defects in *Nckx30c* mutants compared to *w^1118^*controls. We quantified three locomotor parameters for all individuals in the respective groups: the percentage of time spent moving, average speed, and path linearity. **Figure 5C, D** show the changes in these variables in 5-day-old and 30-day-old flies, respectively.

**Figure 5:**
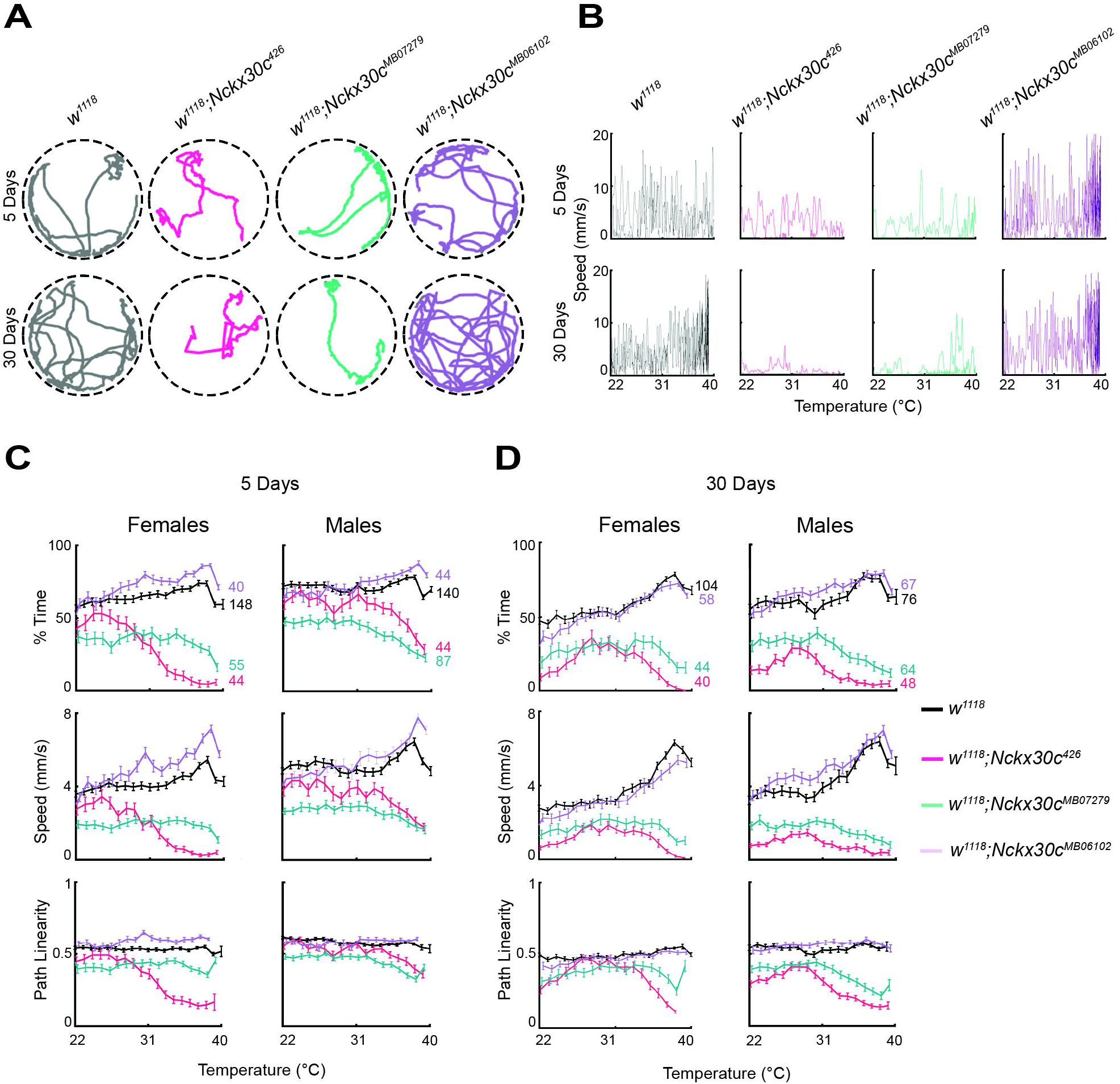
Automated video tracking of locomotor activity in Nckx30c mutants *at* 39°C. **(A)** Representative locomotor trajectories of a single male fly from each genotype recorded at 39°C. Top panel: trace of a 5-day-old fly; bottom panel: trace of a 30-day-old fly. **(B)** Speed profiles of the same representative flies shown in (A) as temperature increased from 22°C to 39°C. For each genotype, the upper trace shows the 5-day-old fly, and the lower trace shows the 30-day-old fly. **(C-D)** Quantification of temperature-dependent locomotor performance for all tested flies. For each genotype, three parameters are plotted as a function of increasing temperature: % time spent moving, average walking speed, and path linearity. Numbers next to the curves in the “% time” plots indicate the sample size for each genotype. **(C)** Data for 5-day-old females and males; **(D)** data for 30-day-old females and males.

At 5 days **(Figure 5C)**, females of both *w^1118^*(n=148) and *w^1118^*;*Nckx30c^MB06102^* (n=40) maintained a similar amount of time throughout the temperature shift, spending approximately 75% and 80% of the time moving at 39°C. In contrast, the percentage of time moving gradually declined for *w^1118^*;*Nckx30c^426^* (n=44) and *w^1118^*;*Nckx30c^MB07279^*(n=55) as the temperature increased from 22°C to 39°C. *w^1118^*;*Nckx30c^426^* spent about 5%, and *w^1118^*;*Nckx30c^MB07279^* about 20%, of the time moving in the arena. A similar pattern was observed in males. *w^1118^* (n=140) spent roughly 70% of the time moving, and *w^1118^*;*Nckx30c^MB06102^*(n=44) about 75% at 39°C.

Meanwhile, *w^1118^*;*Nckx30c^426^*(n=44) and *w^1118^*;*Nckx30c^MB07279^* (n=87) moved approximately 30% and 25% of the time, respectively, at 39°C.

Average speeds of females from *w^1118^*;*Nckx30c^426^* and *w^1118^*;*Nckx30c^MB07279^*gradually decreased with rising temperature, and at 39°C, were approximately 0 mm/s and 1 mm/s at respectively. In comparison, the average speed of *w^1118^* was around 4 mm/s, while *w^1118^*;*Nckx30c^MB06102^* reached about 6 mm/s at 39°C. Similar to females, males also exhibited lower average speeds for *w^1118^*;*Nckx30c^426^* and *w^1118^*;*Nckx30c^MB07279^*, recorded at roughly 2 mm/s at 39°C. Conversely, *w^1118^* and *w^1118^*;*Nckx30c^MB06102^* males showed mean speeds of approximately 4 mm/s and 7 mm/s at the same temperature.

Path linearity for females showed that *w^1118^*;*Nckx30c^426^* was the lowest at approximately 0.2, while *w^1118^*;*Nckx30c^MB07279^* was around 0.4 at 39°C. In contrast, both *w^1118^* and *w^1118^*;*Nckx30c^MB06102^*were about 0.5. Only *w^1118^*;*Nckx30c^426^* exhibited a gradual decline in path linearity with increasing temperature, whereas the other genotypes remained nearly constant.

Male *w^1118^*;*Nckx30c^426^*and *w^1118^*;*Nckx30c^MB07279^* had a path linearity of approximately 0.3 at 39°C, while both *w^1118^* and *w^1118^*;*Nckx30c^MB06102^*consistently maintained around 0.5.

At 30 days of age **(Figure 5D)**, females of both *w^1118^*(n=104) and *w^1118^*;*Nckx30c^MB06102^* (n=58) maintained similar activity levels throughout the temperature shift, spending about 70% of the time moving at 39°C. In contrast, the percentage of time moving was lower for *w^1118^*;*Nckx30c^426^* (n=40) and *w^1118^*;*Nckx30c^MB07279^*(n=44) compared to the other two lines at 39°C. *w^1118^*;*Nckx30c^426^* spent approximately 2% of the time moving, while *w^1118^*;*Nckx30c^MB07279^*spent about 10%. A similar pattern was observed in males, with *w^1118^*(n=76) and *w^1118^*;*Nckx30c^MB06102^* (n=67) both spending roughly 60% of the time moving at 39°C.

Conversely, *w^1118^*;*Nckx30c^426^*(n=48) and *w^1118^*;*Nckx30c^MB07279^* (n=64) moved approximately 5% and 10% of the time at 39°C.

The average speed of females at 30 days old for *w^1118^*;*Nckx30c^426^* and *w^1118^*;*Nckx30c^MB07279^*was approximately 1 mm/s at 39°C. In comparison, both *w^1118^* and *w^1118^*;*Nckx30c^MB06102^* had an average speed of about 5 mm/s at 39°C. Similar to females, male flies of *w^1118^*;*Nckx30c^426^* and *w^1118^*;*Nckx30c^MB07279^* also displayed a lower average speed, around 0 to 1 mm/s, throughout the temperature shift. Conversely, *w^1118^* and *w^1118^*;*Nckx30c^MB06102^* maintained an average speed of approximately 5 mm/s at 39°C.

Path linearity at 30 days for females showed that *w^1118^*;*Nckx30c^426^* had the lowest value, around 0.1, and *w^1118^*;*Nckx30c^MB07279^* was around 0.4 at 39°C. In contrast, both *w^1118^* and *w^1118^*;*Nckx30c^MB06102^* maintained a linearity of about 0.5 throughout the temperature shift. Male flies of *w^1118^*;*Nckx30c^426^* and *w^1118^*;*Nckx30c^MB07279^* exhibited path linearity of approximately 0.2 and 0.3, respectively, at 39°C, while *w^1118^* and *w^1118^*;*Nckx30c^MB06102^* consistently showed a path linearity of about 0.5.

Additionally, to assess the functional consequences of different *Nckx30c* mutations and to explore genetic complexity at this locus, we generated compound heterozygotes of different pairwise combinations between the three alleles: *w^1118^*;*Nckx30c^426^* (point mutation in exon 2), *Nckx30c^MB07279^* (transposable element insertion in exon 2), and *Nckx30c^MB06102^* (transposable element insertion in intron 2) and *w^1118^* controls (**Supplemental Figures S7, S8)**. The combinations were: *w^1118^*;*Nckx30c^426^*^/*+*^, *w^1118^*;*Nckx30c^426^*^/*MB07279*^, *w^1118^*;*Nckx30c^426^*^/*MB06102*^ and *w^1118^*;*Nckx30c^MB07279^*^/*MB06102*^. These heterozygotes were compared against the control *w^1118^*. This approach allowed us to test whether different combinations of *Nckx30c* alleles differentially impact locomotor behavior. We measured the total distance traveled by each compound heterozygote at 5 and 30 days of age for both females and males at 22°C and 39°C. At 22°C (**Supplemental Figure S7)**, 5-day-old female and male *w^1118^*;*Nckx30c^MB07279^*^/*MB06102*^ flies showed significantly reduced locomotion with average traveling distances of 0.11m and 0.17m, respectively, which was approximately 52% and 55% lower than the control *w^1118^* (female: 0.22 m and male: 0.39 m). Whereas for the other allele combinations, females didn’t display any significant change in their traveling distance (*w^1118^*;*Nckx30c^426^*^/*+*^: 0.27 m, *w^1118^*;*Nckx30c^426^*^/*MB07279*^: 0.24 m, and *w^1118^*;*Nckx30c^426^*^/*MB06102*^: 0.31 m). Previously, we observed that homozygous *w^1118^*;*Nckx30c^426^*and *w^1118^*;*Nckx30c^MB07279^* showed a significant reduction in average traveling distance compared to the control. This is also shown by the horizontal dashed lines in **Supplemental Figure S7.** Male *w^1118^*;*Nckx30c^426^*^/*MB07279*^ (0.3 m) and *w^1118^*;*Nckx30c^426^*^/*MB06102*^ (0.38 m) compound heterozygote flies showed a reduction in their average traveling distance by 22% and 2% respectively, compared to *w^1118^* (0.39m).

However, at 30 days, all female and male groups showed a significant reduction in average travel distance compared to the control *w^1118^* (data available in the **Supplemental datasheet 2**). Similar behavior was observed for homozygous *w^1118^*;*Nckx30c^426^*and *w^1118^*;*Nckx30c^MB07279^* (**Supplemental Figure S7,** horizontal dashed lines).

At high temperature (39°C) (**Supplemental Figure S8)**, 5-day-old female and male *w^1118^*;*Nckx30c^426^*^/*MB07279*^ flies displayed an average distance traveled of 0.09m (84% decrease) and 0.06m (90% decrease) compared to control *w^1118^* (female: 0.57m and male: 0.65m). which was the lowest among other compound heterozygotes. Similar behavior was seen for the homozygous *w^1118^*;*Nckx30c^426^* and *w^1118^*;*Nckx30c^MB07279^* (indicated in **Supplemental Figure S8** by the horizontal dashed lines). Although at 39°C we observe a significant reduction of all the other heterozygotes, including *w^1118^*;*Nckx30c^426/MB06102^* and *w^1118^*;*Nckx30c^MB07279/MB06102^*but the homozygous *w^1118^*;*Nckx30c^MB06102^* (shown by the horizontal dashed line in **Supplemental Figure S8**) didn’t have any comparable difference with the control *w^1118^* (data available in the **Supplemental datasheet 2**).

Similarly, both 30-day-old female and male *w^1118^*;*Nckx30c^426^*^/*MB07279*^ flies showed an average distance traveled of about 0.01m (98% decrease) compared to control *w^1118^* (female: 0.60m and male: 0.61m). A similar locomotor pattern was observed in homozygous *w^1118^*;*Nckx30c^426^*and *w^1118^*;*Nckx30c^MB07279^* flies (horizontal dashed lines in **Supplemental Figure S8**). All the other heterozygote combinations also showed a significantly reduced traveling distance, whereas homozygous *w^1118^*;*Nckx30c^MB06102^* didn’t when compared with the control *w^1118^* (data available in the **Supplemental datasheet 2**).

What can be inferred from these results is that the allelic interactions are highly complex in relation to locomotor behavior. At baseline temperature, the locomotor behavior is not just the functional consequence of exon 2 mutations; the intron 2 mutation may also contribute to the phenotype when combined with the other two alleles. We also saw sex-specific differences, which indicate a sex-dependent contribution of *w^1118^*;*Nckx30c^MB06102^* in compound heterozygotes, consistent with what was observed in the *w^1118^*;*Nckx30c^MB06102^* homozygous female flies but at an older age (**Figure 4C**). This implies that although the mutation in *w^1118^*;*Nckx30c^MB06102^*is present in intron 2, the functional consequence of this allele may contribute to the locomotor defect in a heteroallelic combination with other mutant alleles.

### *Nckx30c* mutants exhibit synaptic morphology defects at the larval neuromuscular junction (NMJ) and loss of motor neuron branches in adult dorsal longitudinal muscle (DLM)

Mutations in ion channel genes that disrupt ion homeostasis can lead to morphological defects at the neuromuscular junction (NMJ), often characterized by increased neurotransmitter release and synaptic hyperexcitability (Budnik et al., 1990). We hypothesized that mutations in *Nckx30c* affect synaptic morphology at the larval NMJ. To test this, we analyzed muscle 4 of third-instar larvae, which exhibits stereotypical NMJ morphology and bouton formation (**Figure 6A**). Using immunohistochemistry, we quantified the bouton and satellite bouton numbers from NMJ at muscle 4 from each of the abdominal hemi-segments. In *w^1118^* (n=25), used as a control, we observed 15 ± 3 boutons per NMJ and no satellite boutons. These values served as a baseline for comparison with *Nckx30c* mutant NMJs. Both *w^1118^*;*Nckx30c^426^* and *w^1118^*;*Nckx30c^MB07279^* had 16 ± 2 (*P* = 0.79) and 18 ± 2 (*P* = 0.07) boutons per NMJ, respectively, and the difference was not significant. However, we detected the presence of satellite boutons in both mutants, with *w^1118^*;*Nckx30c^426^* showing 3 ± 2 and *w^1118^*;*Nckx30c^MB0727^* showing 4 ± 2 satellite boutons per NMJ. This represents a significant increase compared to the absence of satellite boutons in the NMJ of *w^1118^* larvae (*P* < 0.0001) for both mutants. The presence of satellite boutons could be associated with abnormal neurotransmitter release, a hallmark of synaptic dysfunction (Rossetto et al., 2011).

**Figure 6:**
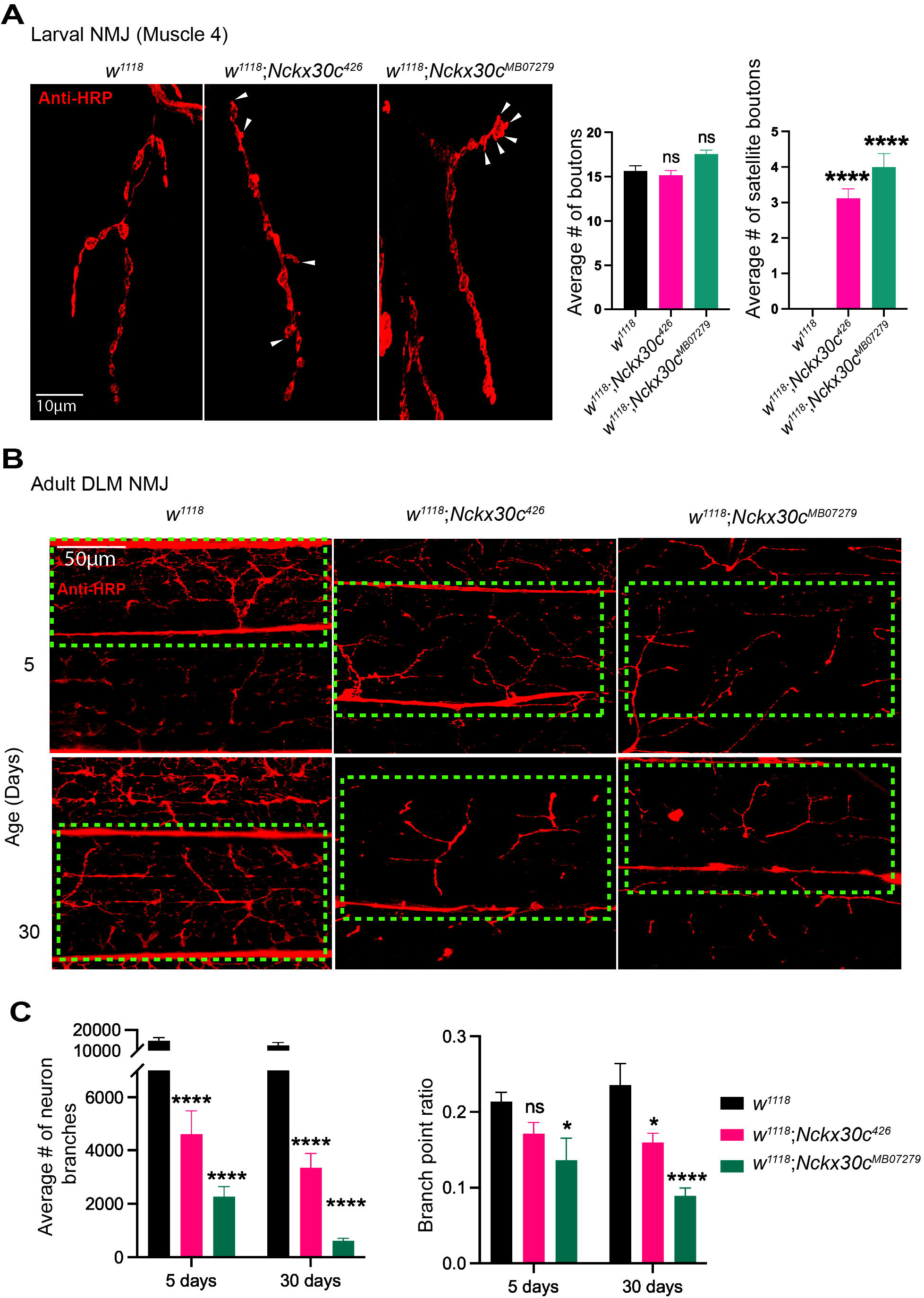
***Nckx30c* mutants show larval and adult (DLM) NMJ defects. (A)** Left panel: Representative confocal images of larval NMJs at muscle 4 stained with Anti-HRP. *w^1118^;Nckx30c^426^* and *w^1118^;Nckx30c^MB07279^* compared with *w^1118^*. White arrowheads indicate satellite boutons at synaptic termini. Right panel: Quantification of bouton numbers and satellite bouton numbers per NMJ is shown by the graph (mean ± SEM). Scale bar:10 μm. For quantification, n = 25 NMJs for *w^1118^*, n = 25 for *w^1118^;Nckx30c^426^* and n = 16 for *w^1118^;Nckx30c^MB07279^*. Each genotype was compared against *w^1118^*.Statistical significance was determined by one-way ANOVA with Dunnett’s post-hoc test; \*\*\*\**P* ≤ 0.0001, *ns* = not significant. Specific P values are reported in the **Supplemental datasheet 1**. **(B)** Representative confocal images of DLM NMJs labeled with Anti-HRP at 60x. Flies were collected at 5 days and 30 days of age for each genotype. For each genotype and age group, n=15-20 thoraces were dissected and stained. Scale bar: 50 μm. The green dotted line marks the boundaries of a single muscle fiber. **(C)** From all the imaged thoraces, only high-quality images with well-preserved muscle fibers were included in the analysis. Left panel: Quantification of the total number of DLM motor neuron branches per hemithorax for each genotype at 5 and 30 days of age (mean ± SEM). Number of individual flies used for this quantification, 5 days: *w^1118^*, n = 5, *w^1118^;Nckx30c^426^*, n = 8 and *w^1118^;Nckx30c^MB07279^*, n = 9; 30 days: *w^1118^*, n = 7, *w^1118^;Nckx30c^426^*, n = 7 and *w^1118^;Nckx30c^MB07279^*, n = 8. Right panel: Branch point ratio for the same individuals as a measure of branching complexity (mean ± SEM). Each genotype was compared against *w^1118^*. Statistical significance was determined by two-way ANOVA with Dunnett’s post-hoc test; Specific P values are reported in the **Supplemental datasheet 1**.

To determine whether the structural integrity of adult neuromuscular junctions (NMJs) is affected in *Nckx30c* mutants, we characterized the motor neuron morphology in dorsal longitudinal muscles (DLMs) in young (5-day-old) and aged (30-day-old) flies (**Figure 6B**).

Adult thoraces were dissected and stained with anti-HRP to visualize motor neuron architecture (Sidisky & Babcock, 2020). 5-day-old *w^1118^* control flies displayed stereotypic highly branched motor neurons innervating the DLM fibers, while both *w^1118^*;*Nckx30c^426^* and *w^1118^*;*Nckx30c^MB0727^* lines exhibited reduced motor neuron branching. By 30 days of age, these defects were exacerbated, with visible loss of axonal branches and reduced innervation of DLM fibers. In contrast, *w^1118^* flies maintained intact with elaborate motor neuron branching. These findings indicate that *Nckx30c* is required to maintain NMJ structure, and its loss leads to progressive reductions in motor neuron branching in DLMs, likely contributing to the locomotor impairments observed in *Nckx30c* mutant flies.

For each genotype we next quantified the average number of neuronal branches at both 5 days and 30 days of age (**Figure 6C**). At 5 days, control *w¹¹¹* (n=5) exhibited an average of 14,682 neuron branches, whereas the *Nckx30c ²* mutants (n=8) showed a significantly lower mean of 4,606 branches (*P* < 0.0001). The *Nckx30c^MB07279^*line (n=9) displayed the most severe reduction, with only 2,270 branches on average (*P* < 0.0001). At 30 days, neuronal branching decreased across all genotypes, but the decline was especially severe in the mutants. *w¹¹¹* (n=7) maintained an average of 12,479 branches, whereas *Nckx30c ²* mutants (n=7) averaged 3,351 branches (*P* < 0.0001). The *Nckx30c^MB07279^* mutants (n=8) exhibited a significant loss, retaining only 622 branches on average (*P* < 0.0001). At both ages, both *Nckx30c* mutant lines showed significantly reduced neuronal branching compared to *w¹¹¹* (p < 0.0001), and this effect became more severe with age.

Branching complexity, expressed as the branch point ratio, was quantified for each genotype at 5 days and 30 days of age (**Figure 6C**). At 5 days, *w¹¹¹* exhibited a branch point ratio of 0.21, while *Nckx30c ²* mutants showed a non-significant decrease to 0.18. In contrast, the *Nckx30c^MB07279^*mutants displayed a significant reduction in branch point ratio to 0.13 (*P* = 0.03). By 30 days, a further decline in branching complexity was observed across genotypes. *w¹¹¹* maintained a ratio of 0.19, whereas *Nckx30c ²* mutants exhibited a significant decrease to 0.16 (*P* = 0.03), and *Nckx30c^MB07279^* mutants showed a significant reduction to 0.08 (*P* < 0.0001).

Together, these results indicate that *Nckx30c* mutations lead to an age-dependent reduction in motor neuron branches and branching complexity, with the *Nckx30c^MB07279^*mutant showing the most severe effect.

### *Nckx30c* mutants display an increased susceptibility to heat-induced seizures and alterations in neural circuit performance

The TS convulsion behavioral phenotype and alterations in NMJ structure in the DLM flight muscles prompted us to assess neural circuit function in adult *Nckx30c* mutant flies. We employed a tethered fly preparation (**Figure 7A**) to monitor spiking of the DLM (Iyengar & Wu, 2014). Several motor circuits drive DLM firing, including grooming and flight pattern generators (Lee et al., 2019), as well as the giant-fiber mediated jump and flight escape reflex (J. E. Engel & Wu, 1996). Additionally, these muscles serve as a convenient readout for seizure activity across the fly nervous system (Iyengar & Wu, 2021; Lee & Wu, 2002; Pavlidis & Tanouye, 1995). We first monitored spontaneously generated firing in *w^1118^*;*Nckx30c^426^*flies at room temperature (∼21°C), comparing them against the *w^1118^*background control flies. We found similar activity patterns in both groups, consisting of occasional spikes which corresponded with grooming activity (**Figure 7B**). There was no significant difference in the overall firing frequency between the respective groups (**Figure 7C**). Notably, none of the *w^1118^*;*Nckx30c^426^*tested flies showed high-frequency spike bursts (>30 Hz) that are characteristic of spontaneous seizure activity in *Drosophila* mutants (e.g. (Landaverde et al., 2024)).

**Figure 7:**
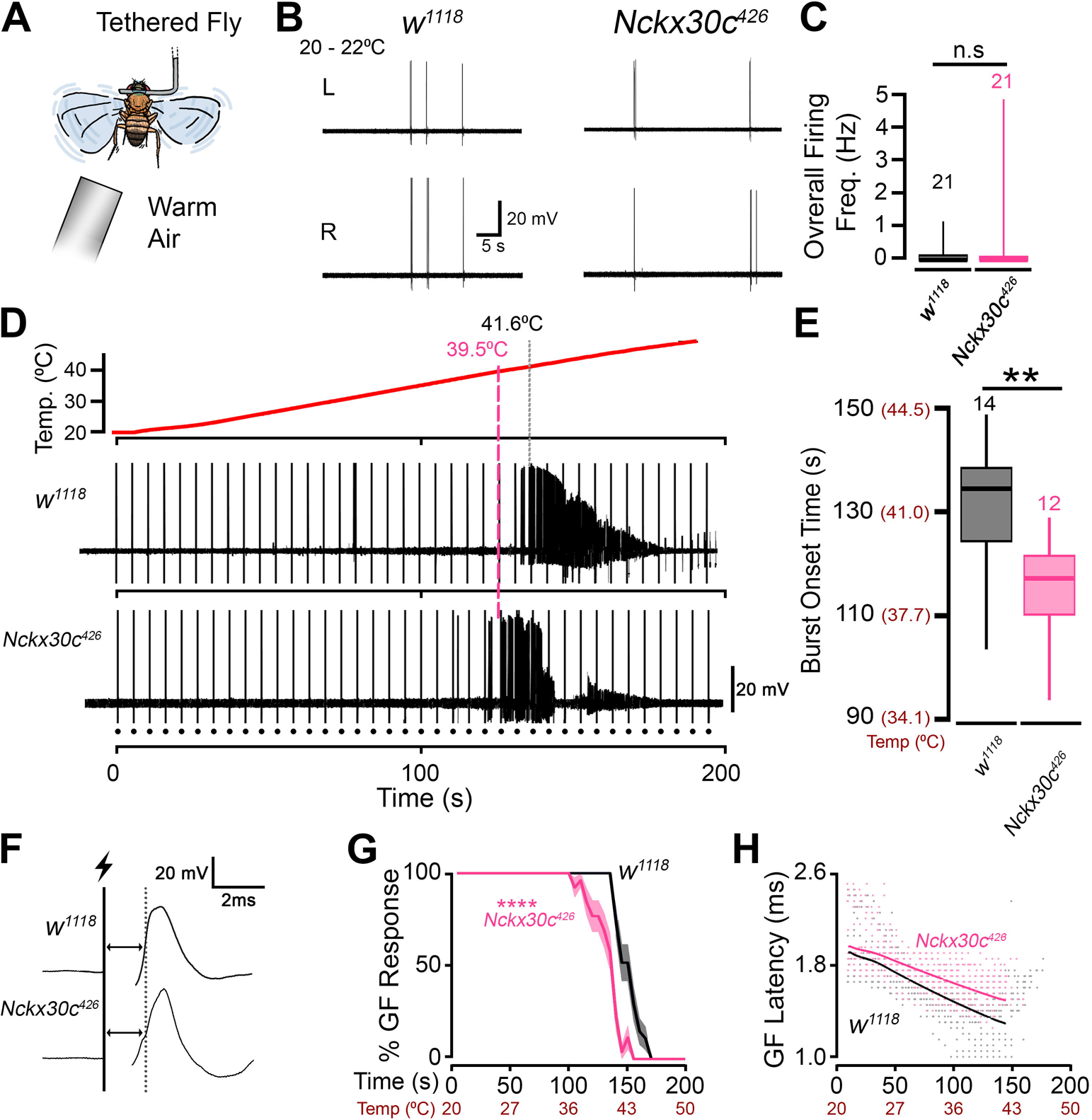
Electrophysiological analysis of motor circuits driving DLM spiking in *Nckx30c* mutants. **(A)** Diagram of the tethered fly preparation. Warm air was blown through a tube (∼3 cm away) to heat the fly. **(B**) Representative traces of DLM spiking activity from the left (L) and right (R) muscles of a *w^1118^* fly and *w^1118^;Nckx30c^426^* mutant at room temperature (20 – 22 °C). **(C)** Box plot of overall firing frequency at room temperature. Numbers above box plots indicate sample size. **(D)** Profile of the temperature ramp protocol and representative traces of activity in a *w^1118^* fly and *w^1118^;Nckx30c^426^* mutant. Onset of DLM spike bursts in the respective flies are marked by dashed lines. Dots indicate artifacts associated with giant fiber stimuli. **(E)** Box plot of burst onset times. Temperatures corresponding to time scale indicated in parentheses. Numbers above box plots indicate sample size. **(F)** Representative traces of giant fiber stimulation-evoked DLM spikes. Lightning bolt indicates electrical stimulation (15 V, 0.1 ms duration). Dotted lines and arrows indicate latency measurement. **(G)** Fraction of stimuli evoking a giant fiber (GF) response during the temperature ramp protocol. Temperatures corresponding with time scale are shown below. Shaded region indicates the standard error of proportion. **(H)** Scatterplot of stimulation-evoked DLM spike latencies during the temperature ramp protocol. Bold lines indicate expected values as a function of time for the distributions based on the linear fits in **Supplemental Figure S9**. Statistical significance in panels **C, E** were determined by rank-sum test, and in panel **G** by a log-rank, \*\**P* ≤ 0.01, \*\*\*\**P* ≤ 0.0001.

The TS convulsion phenotype in *w^1118^*;*Nckx30c^426^*flies (**Figure 5**) prompted us to develop a method of heating the tethered fly on the rig to enable recording of DLM spiking at high temperatures. We used an aquarium pump and process heater to blow gradually warmer air across the fly with a ramp of ∼0.18 °C/s (**Figure 7A**, **D**). As the temperature increased, we monitored the DLM for seizure-related spike bursts. As shown in **Figure 7D, E**, *w^1118^* flies displayed DLM spike bursts indicative of seizures at a median time of 134 s (corresponding to 41.8 °C) during the ramp. In contrast, *w^1118^*;*Nckx30c^426^* flies displayed spiker bursts significantly earlier (median time: 117 s, corresponding to 39.0 °C). These electrophysiological observations are consistent with behavioral observations at 39 °C, where *w^1118^*;*Nckx30c^426^* flies show convulsions while *w^1118^* control flies do not (**Figure 5**).

During the temperature ramp experiments, we generated a series of stimuli (15 V, 0.2 Hz) across the brain to assess performance of the giant fiber escape circuit. These stimuli directly evoke action potentials in the descending giant fiber neurons, which in-turn activates a peripherally synapsing interneuron (PSI) that synapses on the DLM motor neuron, resulting in “short-latency” spike responses in the DLM (J. E. Engel & Wu, 1996) (**Figure** 7**F**). At room temperature, in both *w^1118^;Nckx30c^426^* and *w^1118^*flies, stimulation reliably resulted in a DLM spike response (100%, **Figure 7G**). Furthermore, the response latencies were comparable (*w^1118^;Nckx30c^426^*: 1.95 ± 0.01 ms vs. *w^1118^*: 1.89 ± 0.01 ms), with a slight but statistically significant delay in the response latency of mutants (p = 0.0023). During the temperature ramp, obvious differences between the groups arose. As shown in **Figure 7G**, *w^1118^;Nckx30c^426^* flies, stimuli failed to evoke responses earlier (median: 140 s) compared to *w^1118^* counterparts (median: 155 s). Notable differences were also evident in scatterplots of the DLM latencies as a function of time and temperature (**Figure 7H**). Before failing at high temperatures, *w^1118^* flies displayed shorter spike latencies compared to room temperature (**Figure 7H**, e.g. 1.32 ms at 110 s). In *w^1118^;Nckx30c^426^* flies the magnitude of the latency decrease was markedly smaller (1.62 ms at 110 s). Indeed, when the temperature was plotted against response latency on a semi-log scale, linear fits of the data indicated a significantly steeper temperature dependence coefficient in *w^1118^* flies compared to *w^1118^;Nckx30c^426^* mutants (**Supplemental Figure S9,** p = 3.73 x 10^-10^). Together, our findings indicate clear *Nckx30c* mutation-related alterations in temperature-dependent characteristics of neurotransmission along the giant fiber circuit.

### Neuron-specific knockdown of *Nckx30c* results in TS convulsions, shorter lifespan, and locomotor defects, while glial knockdown causes age-dependent locomotor decline

To determine the cell-type specificity of *Nckx30c* function underlying the observed TS convulsion phenotype, we performed targeted knockdown experiments in the nervous system (**Figure 8A**). Notably, previous studies have shown that the NCKX family member *zydeco* induces TS paralysis and Ca² fluctuations when knocked down specifically in glial cells (Melom & Littleton, 2013), prompting us to assess whether *Nckx30c* has a similar or distinct cellular requirement.

**Figure 8:**
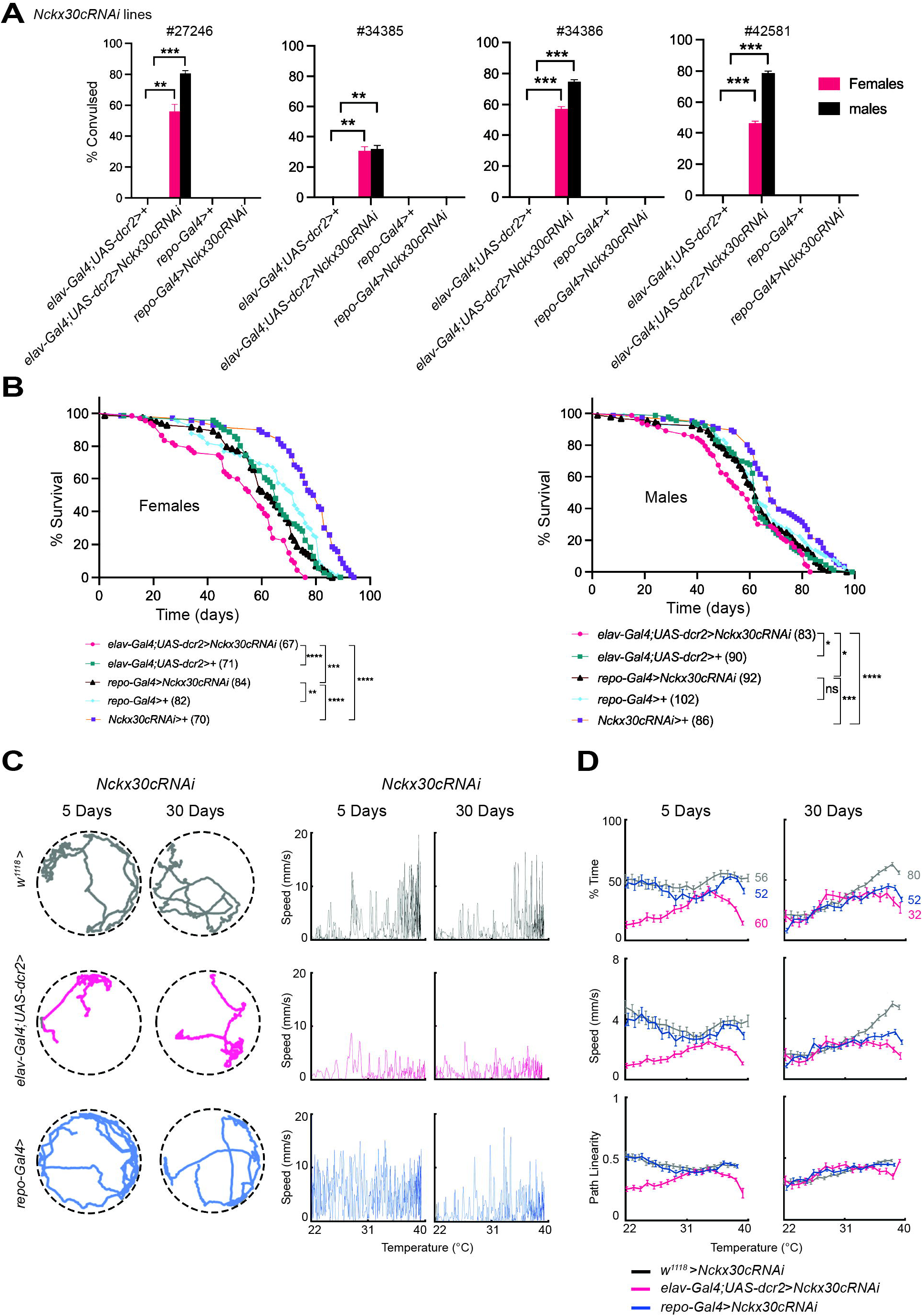
Knockdown of *Nckx30c* in neurons and glia differentially affects behavioral phenotypes and lifespan. **(A)** TS paralysis assay of *Nckx30c* knocked down in neurons (*elav-Gal4;UAS-dcr2>UAS-Nckx30cRNAi*) and glia (*repo-Gal4>UAS-Nckx30cRNAi*). Four different *Nckx30cRNAi* lines are used (#27246, #34385, #34386, and #42581). For each genotype, the sample size is n = 75. Values shown are mean ± SEM; each knockdown genotypes are compared against its control and statistical significance was determined by using One sample t and Wilcoxon test; \*\**P* ≤ 0.01, \*\*\**P* ≤ 0.001, P values are reported in the **Supplemental datasheet 1. (B)** Lifespan curves for flies of the indicated genotypes maintained at 25°C. Numbers in parentheses indicate sample size. Log-Rank (Mantel-Cox); \**P* ≤ 0.05, \*\**P* ≤ 0.01, \*\*\**P* ≤ 0.001, \*\*\*\**P* ≤ 0.0001, *ns* = not significant. P values are reported in the **Supplemental datasheet 1. (C)** Left panel: Representative locomotor trajectories of a single female from each knockdown genotype recorded at 39°C, shown for both 5-day and 30-day-old flies. Right panel: speed profiles of the same representative flies as temperature increased from 22°C to 39°C. **(D)** Quantification of temperature-dependent locomotor performance for all tested flies. For each knockdown genotype, three parameters are plotted as a function of increasing temperature: % time spent moving, average walking speed, and path linearity. Numbers next to the curves in the “% time” plots indicate the sample size for each knockdown genotype.

We used a *UAS-Nckx30cRNAi* line (#27246) to reduce *Nckx30c* expression in a cell-type-specific manner in the nervous system and tested TS convulsions **(Figure 8A)**. Neuronal knockdown was achieved using *elav-Gal4*;*UAS-dcr2,* and glial knockdown using *repo-Gal4*.

Flies with neuronal knockdown, *elav-Gal4*;*UAS-dcr2>UAS-Nckx30cRNAi* (n=75), showed TS convulsion, with 42 out of 75 (56%) of female and 60 out of 75 (80%) of male flies displaying the phenotype. In contrast, the glial knockdown, *repo-Gal4>UAS-Nckx30cRNAi* (n=75), flies didn’t show any convulsion for either sex. To validate the specificity of the convulsive phenotype, we tested three additional *UAS-Nckx30cRNAi* lines (#34385, #34386, and #42581). All RNAi lines targeting neurons using *elav-Gal4;UAS-dcr2* recapitulated the TS convulsive phenotype. With *UAS-Nckx30cRNAi* (#34385), 23 out of 75 (∼31%) females and 24 out of 75 (32%) males; *UAS-Nckx30cRNAi* (#34386), 42 out of 75 (∼56%) females and 55 out of 75 (74%) males and with *UAS-Nckx30cRNAi* (#42581), 33 out of 75 (∼44%) females and 58 out of 75 (78%) males displayed the TS phenotype. For knockdown experiments, the genetic controls were: *elav-Gal4;UAS-dcr2>*+ (driver control), *repo-Gal4>*+ (driver control), and *UAS-Nckx30cRNAi>*+ (RNAi control).

For our subsequent *UAS-Nckx30c-RNAi* experiments, we used line #27246. To verify knockdown efficiency, we performed RT-qPCR (**Supplemental Figure S10**) and found that *elav-Gal4*;*UAS-dcr2>UAS-Nckx30cRNAi* flies showed a reduction in *Nckx30c* expression in whole heads compared to the control *elav-Gal4;UAS-dcr2>*+, confirming knockdown was achieved. We also observed a reduction in *Nckx30c* expression in *repo-Gal4>UAS-Nckx30cRNAi* flies compared to their control *repo-Gal4>*+.

Next, we assessed the effect of *Nckx30c* knockdown on lifespan using *elav-Gal4;UAS-dcr2>UAS-Nckx30cRNAi* and *repo-Gal4>UAS-Nckx30cRNAi* (**Figure 8B**). In females, neuronal knockdown *elav-Gal4;UAS-dcr2>UAS-Nckx30cRNAi* (n=67) significantly reduced lifespan compared to both controls *elav-Gal4;UAS-dcr2>*+ (n=71); *P* <0.0001 and *UAS-Nckx30cRNAi>*+ (n=70); *P* <0.0001, and almost all the flies died before 80 days. Glial knockdown flies, *repo-Gal4>UAS-Nckx30cRNAi* (n=84) also showed reduced lifespan compared to control *repo-Gal4>*+ (n=82); *P* =0.0086 and *UAS-Nckx30cRNAi>*+ (n=70); *P* <0.0001, where almost all the flies died before 90 days. Furthermore, neuronal knockdown flies *elav-Gal4;UAS-dcr2>UAS-Nckx30cRNAi* showed significantly shorter lifespan compared to glial knockdown flies *repo-Gal4>UAS-Nckx30cRNAi* (*P* =0.0006). Similarly, *elav-Gal4;UAS-dcr2>UAS-Nckx30cRNAi* (n=83) males displayed significantly reduced lifespan compared to both controls *elav-Gal4;UAS-dcr2>*+ (n=90); *P* =0.0260 and *UAS-Nckx30cRNAi>*+ (n=86); *P* <0.0001. The difference in lifespan was not as pronounced as observed in the females. *repo-Gal4>UAS-Nckx30cRNAi* (n=92) didn’t display any significant change in their lifespan compared to control *repo-Gal4>*+ (n=102); but they showed significantly shorter lifespan than *UAS-Nckx30cRNAi>*+ (n=86); *P* =0.0004. Compared to glial knockdown flies, *repo-Gal4>UAS-Nckx30cRNAi*, neuronal knockdown flies, *elav-Gal4;UAS-dcr2>UAS-Nckx30cRNAi,* showed significantly shorter lifespan; *P* =0.0168. From the observed results, we found that for both females and males, neuron-specific *Nckx30c* knock down resulted in the shortest lifespan among all the genotypes examined. These findings collectively indicate that *Nckx30c* plays a role in neuronal function, where its knockdown is sufficient to induce TS convulsions, early-onset locomotor defects, and a significantly shortened lifespan, while glial knockdown contributes to locomotor decline at later ages and reduces lifespan relative to controls.

Finally, we examined locomotor activity in *Nckx30c* knockdown flies using the automated video tracking system at 39°C (**Figure 8C, D**). Assays were performed on both 5-day and 30-day-old female flies, across *elav-Gal4*;*UAS-dcr2>UAS-Nckx30cRNAi* and *repo-Gal4>UAS-Nckx30cRNAi*, compared to the RNAi control *UAS-Nckx30cRNAi>+* (where *UAS-Nckx30cRNAi* crossed with w^1118^). Representative movement tracks of a single female fly at 39°C (**Figure 8C**) show the locomotor defects in *elav-Gal4*;*UAS-dcr2>UAS-Nckx30cRNAi* flies at 5 days, and in both *elav-Gal4*;*UAS-dcr2>UAS-Nckx30cRNAi* and *repo-Gal4>UAS-Nckx30cRNAi* flies at 30 days, relative to the control.

To further characterize this impairment, we quantified the speed of the same representative flies as temperature increased from 22°C to 39°C (**Figure 8C**). At 5 days, *elav-Gal4*;*UAS-dcr2>UAS-Nckx30cRNAi* reached a maximum speed of ∼4 mm/s whereas the control *UAS-Nckx30cRNAi>+* reached approximately 15 mm/s. *repo-Gal4>UAS-Nckx30cRNAi* wasn’t much different than the control, having a speed of ∼14 mm/s. However, at 30 days, both *elav-Gal4*;*UAS-dcr2>UAS-Nckx30cRNAi* and *repo-Gal4>UAS-Nckx30cRNAi* showed reduced speed of ∼3 mm/s and ∼9 mm/s respectively compared to the control.

We then quantified three locomotor parameters for all female individuals in each knockdown genotype during the temperature shift from 22°C to 39°C. The variables included the percentage of time spent moving, average speed, and path linearity at both 5 days and 30 days of age (**Figure 8D**). At 5 days, both *repo-Gal4>UAS-Nckx30cRNAi* (n=52) and the control *UAS-Nckx30cRNAi>+* (n=56) maintained comparable activity at 39°C, spending approximately 40% and 50% of the time moving, respectively. In contrast, *elav-Gal4*;*UAS-dcr2* > *UAS-Nckx30cRNAi* females (n=60) showed impaired locomotion, moving only ∼10% of the time at 39°C. However, at 30 days, we see a decline in movement in the knockdown lines. Both *elav-Gal4*;*UAS-dcr2>UAS-Nckx30cRNAi* (n=32) and *repo-Gal4>UAS-Nckx30cRNAi* (n=52) spent approximately 25% of the time moving at 39°C compared to the control *UAS-Nckx30cRNAi>+* (n=80), whose moving time was ∼65%.

The average speed profiles were similar to the patterns observed for the percentage of time spent moving. *elav-Gal4;UAS-dcr2>UAS-Nckx30cRNAi* females showed the most severe impairment, reaching only ∼1 mm/s at 39°C at both 5 days and 30 days of age. In comparison, *repo-Gal4>UAS-Nckx30cRNAi* and the genetic control *UAS-Nckx30cRNAi*>+ showed higher speeds of approximately 3 mm/s and 4 mm/s, respectively, at 5 days. By 30 days, the average speed of *repo-Gal4>UAS-Nckx30cRNAi* further declined to ∼2 mm/s, whereas *UAS-Nckx30cRNAi*>+ females maintained relatively robust locomotion with speeds of ∼5 mm/s at 39°C.

Path linearity for females showed that at 5 days, *elav-Gal4;UAS-dcr2>UAS-Nckx30cRNAi* was approximately 0.2, whereas both *repo-Gal4>UAS-Nckx30cRNAi* and *UAS-Nckx30cRNAi*>+ showed a path linearity of roughly 0.5. On the other hand, all the knockdown genotypes showed a path linearity of ∼0.5 at 30 days of age at 39°C.

These results indicate that neuronal knockdown of *Nckx30c* leads to TS convulsions, shorter lifespan, and early-onset locomotor defects. In comparison, glial knockdown produces a later-appearing, age-dependent locomotor impairment with shorter lifespan.

## Discussion

This study identifies *Nckx30c* as a potential regulator of neuronal excitability, synaptic integrity, and proper function of the nervous system in *Drosophila melanogaster*. Through a combination of forward genetics and complementation testing analysis, we identified a loss-of-function *Nckx30c* mutant that displays temperature-sensitive convulsions, age-dependent neurodegeneration, progressive locomotor decline, and shortened lifespan. In *426* mutant flies, we mapped the mutation to exon 2 of the *Nckx30c* gene within its conserved Na^+^/Ca^2+^ exchange domain. This highlights what is likely to be a functional disruption of the K^+^-dependent Na^+^/Ca^2+^ exchanger encoded by *Nckx30c*.

Temperature sensitive (TS) paralysis in *Drosophila* has been traditionally utilized to identify genes that are crucial for neuronal excitability, including ion channels and Ca^2+^ regulators, revealing key proteins in ion conductance (Ganetzky & Wu, 1986; Suzuki et al., 1971). A major advantage of forward TS screens is that they allow unbiased identification of genes required for neuronal signaling simply by observing behavioral paralysis, without prior knowledge of the underlying gene or protein. This screening technique is also useful in identifying mutations that display convulsion or seizure-like behavior (P. Wang et al., 2004). The TS convulsive phenotype observed in *Nckx30c^426^* mutants aligns with previous reports that associate TS paralysis with impaired neuronal excitability and ion homeostasis (Palladino et al., 2002; Suzuki et al., 1971). Given its proposed role as a Ca^2+^ exchanger, our findings suggest that *Nckx30c* may contribute to maintaining neuronal Ca^2+^ homeostasis under high temperature stress. Interestingly, the transposable element insertion mutant *Nckx30c^MB07279^*, which disrupts the same exon as the point mutant, also displayed TS convulsion and reduced *Nckx30c* gene expression, whereas *Nckx30c^MB06102^*, with an insertion in an intron, did not. This suggests that mutations directly affecting coding regions or regulatory exons of *Nckx30c* are necessary to elicit the phenotype, potentially due to a more severe loss of function.

A hallmark feature of many TS mutants is their association with progressive neurodegeneration (Palladino et al., 2002). Consistently, both *Nckx30c^426^* and *Nckx30c^MB07279^* mutants developed age-dependent brain neuropathology, accompanied by a significantly shortened lifespan. These results reinforce the emerging view that ionic imbalances, particularly calcium dysregulation, are a key contributor to neuronal vulnerability with age (Gleichmann & Mattson, 2011; Stolwijk et al., 2016). The observation that neurodegeneration becomes more pronounced at older age may reflect cumulative mitochondrial dysfunction and excitotoxic stress, possibly arising from sustained calcium imbalance. Although we did not test whether seizure activity accelerates this degeneration, it remains a possibility because recurrent seizures in humans are known to worsen neuronal vulnerability and cognitive decline in epileptic disorders (Klein et al., 2025). Future studies examining whether induced or spontaneous convulsions intensify age-related neuropathology in *Nckx30c* mutants will help clarify the interplay between Ca^2+^ extrusion defects, excitability-dependent stress, and neuronal survival.

Studies of vertebrate retinal NCKX suggest that NCKX family members can form homodimers. It was shown that both NCKX1 and NCKX2 exist as multimeric complexes in the plasma membrane of eye photoreceptor cells (Kang et al., 2003; Schwarzer et al., 1997). Whether this feature of NCKX is also present in cells of the nervous system has not yet been determined (Al-Khannaq & Lytton, 2022). As the protein encoded by *Nckx30c* shares 66% and 71% sequence similarity with NCKX1 and NCKX2, respectively (Haug-Collet et al., 1999), and is a functional homolog of NCKX, it may also form a multimeric complex. In this context, complementation patterns were observed by constructing compound heterozygotes with different *Nckx30c* alleles in our study. The locomotor analysis using compound heterozygotes revealed that only *Nckx30c^426^*/*^MB07279^* exhibited a significant defect in movement, particularly under high-temperature conditions, while other allele combinations did not. This finding suggests that the two exon 2 mutations, *Nckx30c^426^* (point mutation) and *Nckx30c^MB07279^*(transposable element insertion), represent loss-of-function alleles that fail to complement one another. Their combined effect likely disrupts *Nckx30c* function below the threshold required to maintain regular neuronal activity and locomotor behavior. In contrast, we found that heterozygotes involving *Nckx30c^MB06102^* (intronic transposable element insertion) exhibited behavior comparable to *w^1118^*;*Nckx30c^426^*^/*+*^ heterozygotes, indicating that this allele may have minimal or no functional impact on gene expression. The absence of locomotor impairment in these combinations highlights the importance of exon integrity for *Nckx30c* function. The locomotor defect observed in *w^1118^*;*Nckx30c^426/MB07279^* flies was most noticeable under high-temperature stress, and the severity increased with age. These findings suggest that *Nckx30c* plays a role in maintaining neuronal integrity and locomotor function under conditions of physiological stress. However, we note that our data are based on genetic analysis and do not directly demonstrate Nckx30c oligomerization in *Drosophila*.

Over the years, *Drosophila* has enabled the identification and functional characterization of numerous genes and pathways involved in neuroprotection and neurodegeneration, many of which are conserved in humans (Bilen & Bonini, 2005). Additionally, studies of the *Drosophila* larval neuromuscular junction (NMJ), a synaptic link between the motor nerve’s terminal end and a muscle, have provided key insights into the mechanisms regulating synapse development, growth, and neurotransmission. Functionally, these glutamatergic synapses resemble mammalian central synapses and undergo structural remodeling in response to neuronal activity (Collins & DiAntonio, 2007; Gramates & Budnik, 1999; Juel, 2012). The larval NMJ is the classic model system for studying ion channels (such as K^+^, Ca^2+^), mutants for which often show altered bouton number and synaptic overgrowth with satellite boutons (Lee & Wu, 2010; Menon et al., 2013). The adult dorsal longitudinal muscle (DLM) NMJ, an indirect flight-muscle synapse essential for escape, provides a complementary aging model. Its stereotyped innervation permits precise quantification of motor-neuron branching and connectivity (Allen et al., 2006). In *mayday* and *shibire* fly mutants, DLM fibers show progressive, age-dependent loss of motor neuron branching (Beramendi et al., 2007; Sidisky & Babcock, 2020; Sidisky et al., 2021). Studying NMJ integrity is essential for identifying molecular factors underlying age-related synaptic decline and neurodegeneration (Pratt et al., 2021). Our larval NMJ analysis revealed a significant synaptic abnormality in *Nckx30c* mutants with the appearance of satellite boutons.

This feature is often associated with elevated synaptic activity and altered neurotransmission (Budnik et al., 1990; Rossetto et al., 2011), suggesting that loss of *Nckx30c* disrupts synaptic structure and function, possibly due to persistent Ca^2+^ elevation in presynaptic terminals. Our analysis of DLM neuromuscular junctions reveals synaptic abnormalities in *Nckx30c* mutants, indicating that *Nckx30c* is required for proper DLM NMJ morphology and stability, but our data do not distinguish between developmental and adult-specific roles. The early reduction in motor neuron branching observed in both *Nckx30c^426^* and *Nckx30c^MB07279^* mutants, which becomes more pronounced with age, suggests that *Nckx30c* is required not only at early development but also for the long-term stability of NMJs. This progressive denervation is consistent with the locomotor impairments seen in mutant flies and could represent a possible link between disrupted Ca^2+^ extrusion and synaptic maintenance (Vonhoff & Keshishian, 2017). Similar age-dependent NMJ degeneration has been reported in other models of neuronal dysfunction (Sidisky & Babcock, 2020), further supporting the idea that Ca^2+^ homeostasis is essential for preserving motor neuron integrity. Thus, *Nckx30c* may function as a neuroprotective factor, with its loss predisposing flies to both structural synaptic decline and associated behavioral deficits.

To provide functional analysis of the DLMs in adult *Nckx30c* mutants, we employed a tethered fly preparation in which muscle spikes are monitored in intact and behaving flies.

Previous studies have employed similar systems to characterize NMJ and motor circuit function in several excitability mutants (e.g. *Slowpoke* (Elkins et al., 1986), *Shaker* (Jeff E. Engel & Wu, 1998), and *quiver* (Ruan et al., 2017)) including models of epilepsy (*e.g. bss* (Lee & Wu, 2002), and *sugarlethal* (Chi et al., 2019)) as well as mutants prone to neurodegeneration (e.g. *sod1* (Iyengar et al., 2022)). Our initial observations in *Nckx30c^426^* flies did not reveal clear distinctions in spiking patterns from control flies, despite the apparent structural differences at the DLM NMJ. This finding may not be entirely surprising given several instances in larval abdominal NMJs where obvious structural differences (e.g. satellite boutons or excess branching) do not always correlate with alterations in basic neurotransmission characteristics. Consistently, at room temperature, performance of the giant fiber jump-and-flight escape circuit was largely similar between *Nckx30c* mutants and control flies. This relatively simple and well-characterized escape circuit, consisting of the giant fiber neuron, a peripherally synapsing interneuron, DLM motor neuron, serves as a convenient assessment of action potential generation and propagation, as well as neurotransmission across cholinergic and electrical synapses (Allen et al., 2006).

To uncover potential temperature-dependent electrophysiological phenotypes in tethered flies, we developed a new system to blow progressively warmer air on the fly. Similar heating approaches have previously been employed in restrained fly preparations to monitor DLM activity (Kroll et al., 2015). The spike bursts elicited at ∼ 38-40 °C in *Nckx30c* mutants indicate clear vulnerability to heat-evoked seizures consistent with behavioral observations. Bursting in *w^1118^* manifest at a higher temperature range than *Nckx30c* mutants ∼ 40-42 °C, that was consistent with maximal thermal tolerances of wild type *Drosophila* (Folk et al., 2007).

Conceivably, alterations in human NCKX properties could contribute to seizures triggered at high temperatures (e.g. during fever) a well-known phenomenon in the context of several epilepsy syndromes (Feng & Chen, 2016; Verbeek et al., 2015). During temperature ramp experiments, we continuously stimulated the giant fiber pathway to assess its performance.

Although the latency of both *w^1118^* flies and *Nckx30c* mutant responses were faster at higher temperatures (prior to failure), the temperature dependence of this effect was significantly reduced in *Nckx30c* mutants. Given the robust transmission observed at room temperature, a potential implication of these findings is NCKX function acts to ensure robust synaptic transmission performance across a range of environmental temperatures.

Tissue-specific RNAi experiments demonstrated that *Nckx30c* is required in neurons for the temperature-sensitive (TS) convulsion phenotype, as this phenotype was observed only with neuronal knockdown and not with glial knockdown. Locomotor behavior analyses revealed that at a young age (5-day-old), defects were restricted to flies with neuronal *Nckx30c* knockdown. However, at an older age (30-day-old), glial knockdown flies also exhibited locomotor impairments. Lifespan assays further showed that neuronal knockdown resulted in the shortest lifespan, while glial knockdown also led to a significantly reduced lifespan compared to both glial Gal-4 driver and RNAi controls. The age-dependent locomotor defects and reduced lifespan associated with glial knockdown suggest that *Nckx30c* function in glial cells becomes increasingly important with age, thereby contributing to long-term neuronal health. Notably, the other NCKX family member *zydeco,* whose TS phenotype is glia-dependent (Melom & Littleton, 2013), suggests that *Nckx30c* and *zydeco* serve distinct cellular roles in TS convulsions, but may share overlapping functions related to glial contributions in aging.

Mutations in *Nckx30c* could potentially affect the expression of other genes involved in calcium homeostasis. In that regard, we assessed the mRNA levels of *MCU* (Mitochondrial Calcium Uniporter), involved in Ca^2+^ transport into the mitochondria (Choi et al., 2017) and *PMCA* (Plasma Membrane Calcium ATPase) which plays a role in Ca^2+^ extrusion (Erhardt et al., 2021), in 7 days-old male heads of all the *Nckx30c* mutants investigated in this study, but did not observe any significant changes compared to the control strain (**Supplemental Figure S11**). This suggests that the phenotypes observed at 7 days of age in male flies are linked to *Nckx30c* dysfunction rather than secondary alterations in other Ca^2+^ transport mechanisms.

However, analysis of these two candidates does not exclude the possibility that other Ca^2+^ transporters or signaling components are differentially regulated in *Nckx30c* mutants.

Conversely, we did not check gene expression at an older age (30-day-old) when there could have been a compensatory effect at later time points. Also, we cannot exclude that such changes occur in discrete cells or subtypes of cells in the nervous system. Future experiments could address this by using *in-situ* hybridization or immunostaining to visualize and quantify mRNA and/or protein corresponding to these genes in dissected brains of mutants and compare them to controls. It is also possible that the reduced longevity, locomotor defects, and neurodegeneration associated with *Nckx30c* disruption may be the result of mitochondrial damage due to Ca^2+^ overload or oxidative stress. While *Drosophila* mitochondria are capable of undergoing permeability transition (von Stockum et al., 2011) they are resistant to Ca^2+^ overload (Chaudhuri et al., 2016). Mitochondrial permeability transition refers to the opening of a large, non-selective pore in the inner mitochondrial membrane in response to calcium overload and oxidative stress (Carraro & Bernardi, 2023). This event collapses the mitochondrial membrane potential, disrupts ATP production, and can trigger cell death pathways (Bernardi et al., 2006). Given that *Drosophila* mitochondria can undergo permeability transition (von Stockum et al., 2011), it is possible that impaired Ca^2+^ clearance arising from Nckx30c disruption could lead to excessive cytosolic and mitochondrial Ca^2+^ accumulation, thereby promoting mitochondrial permeability transition. This provides a plausible mechanistic link between Ca² dysregulation, mitochondrial dysfunction, and the reduced lifespan, locomotor defects, and neurodegeneration observed in *Nckx30c* mutants. In mammalian cells, the *PPIF* gene (encoding Cyclophilin D) is a critical regulator of permeability transition (Baines et al., 2005) that is inhibited by cyclosporin A. While *Drosophila* mitochondrial permeability transition is insensitive to cyclosporin A (von Stockum et al., 2011), knockdown of the putative *Drosophila* homolog of Cyclophilin D, Cyclophilin 1 (encoded by *Cyp1*), does inhibit permeability transition in cultured *Drosophila* S2R+ cells (Chaudhuri et al., 2016). Additionally, deletion of *Cyp1* is protective against oxidative stress and has been shown to improve lifespan and locomotor defects in a *Drosophila DJ-1*β mutant line that is used as a model of Parkinson’s disease (Kim et al., 2018). Thus, mechanistic insight may be gained by future studies designed to examine whether *Cyp1* deletion could similarly recover the defects associated with *Nckx30c* disruption.

Looking ahead, several experiments could extend our mechanistic understanding. Genetic rescue approaches also hold promise: reintroducing *Nckx30c* itself or expressing K leak channels could help determine whether the observed mutant phenotypes arise solely from Ca^2+^ accumulation or whether altered K^+^ handling also contributes. Overexpressing Ca^2+^ leak channels could work as an attempt to rescue the mutant phenotype. Finally, exploring genetic interactions between *Nckx30c* mutants and models of neurodegeneration such as AD models could provide insight into how Ca^2+^ dysregulation intersects with amyloid-driven pathology. While the connection between NCKX family members and human NDs has not been extensively investigated, the human *Nckx30c* ortholog *SLC24A2* (NCKX2) is suggested to play a role in neuronal Ca^2+^ extrusion and has been implicated in NDs, more recently, in AD. One study based on genome-wide association using whole-genome sequencing of apolipoprotein E ε4 (APOE ε4) carriers identified *SLC24A2* as one of twelve loci significantly associated with AD risk. Here 500 individual participated in the study with 331 of them being AD patients (Park et al., 2021).

Another study reported an elevated levels of NCKX2 phospho-peptides in the serum of AD patients compared to controls, although it remains uncertain whether this increase reflects NCKX2 release from defective neurons or changes in the expression of NCKX2 and post-translational modifications (Florentinus-Mefailoski et al., 2021). While this study only examined n= 24 individuals among whom 12 were AD patients, these findings suggest that disruption of NCKX2 function may contribute to neurodegenerative vulnerability. Testing if defective *Nckx30c* intensifies phenotypes would support the idea that defective ionic balance is an upstream vulnerability factor in amyloid-mediated disease.

Together, these future directions will not only clarify the neuroprotective role of *Nckx30c* but also establish its broader relevance to seizure susceptibility and neurodegenerative diseases.

## Conclusion

This study provides novel insights into the role of the *Drosophila* gene *Nckx30c*, a member of the NCKX family of K-dependent Na^+^/Ca^2+^ exchangers, in regulating neuronal excitability, synaptic integrity, and organismal health. Through a forward genetic screen, a novel mutant allele (*Nckx30c^426^*) was identified that exhibits temperature-sensitive convulsions, age-dependent neurodegeneration, and locomotor dysfunction. Analysis of additional alleles including *Nckx30c^MB07279^*, revealed that mutations disrupting exon 2 are particularly deleterious, emphasizing the importance of this region for ion exchanger function.

Phenotypic characterization across flies from young and old cohorts established a consistent pattern of dysfunction, where the mutant phenotype gained severity with age. Mutants displayed behavioral abnormalities at increased temperature, reduced lifespan, progressive neurodegeneration, and structural synaptic defects at both larval and adult neuromuscular junctions. Specifically, the presence of satellite boutons at larval NMJs and progressive branch loss in adult DLM NMJs revealed that *Nckx30c* may have a role in maintaining synaptic stability throughout life. Tissue-specific knockdown experiments further demonstrated that neuronal *Nckx30c* is essential for preventing temperature sensitive convulsions, early onset of locomotor defect and maintaining lifespan, while glial knockdown contributes to age-dependent locomotor decline.

Together, our findings establish *Nckx30c* as an important regulator of Ca^2+^-dependent neuronal processes in *Drosophila melanogaster*. By linking impaired Ca^2+^ extrusion to seizure susceptibility, synaptic abnormalities, and neurodegeneration, this work highlights a unifying mechanism through which ionic imbalance can trigger both acute and progressive neural dysfunction. This work broadens our understanding of the NCKX family by moving beyond the well-studied glial exchanger *zydeco* and uncovering *Nckx30c*, a counterpart that functions at an early age in neurons and exerts effects in glia, particularly at older ages.

The broader implications of this study are twofold. First, it reinforces the concept that Ca^2+^ dysregulation is a central mechanism connecting seizures and neurodegenerative diseases. Second, it positions *Nckx30c* as a valuable genetic model for exploring how disruptions in Ca^2+^ transport contribute to neurodegenerative disorders and epileptic seizures. By integrating genetic, behavioral, and structural approaches, this study lays the foundation for future electrophysiological, and disease-interaction studies that will further define the molecular pathways linking Ca^2+^ homeostasis, synaptic health, and neuronal survival.

In conclusion, the work presented here advances our understanding of Ca^2+^ regulation in the nervous system by establishing *Nckx30c* as a key factor in neuronal excitability and synaptic maintenance. It also provides a framework for future research into the role of NCKX exchangers in neurodegenerative disease and seizure susceptibility, thereby contributing to the broader goal of uncovering mechanisms that safeguard neuronal function across the lifespan.

## Acknowledgements

We thank Dr. Barry Ganetzky in whose laboratory this work started originally; Dr. Steven Robinow for providing the collection of ENU-mutagenized stocks used in the genetic screen; Luke Chandler for help with the forward genetic screen; Katy Marotto for help with histology, Dr. Kim Lackey at the Optical Analysis Facility at the University of Alabama for assistance with confocal microscopy. We are grateful to all our colleagues in the Chtarbanova and Iyengar labs and members of the Ganetzky joint lab meeting for advice and helpful discussions about the project.

## Data availability

Material including supplemental figures, tables, and datasheets that support the findings of this study are available in the Open Science Framework (OSF) repository at https://osf.io/yqdvm/ (https://doi.org/10.17605/OSF.IO/YQDVM) (Majlish et al., 2025).

## Authors contributions

SC conceptualized the study with input from AI and ANKM; ANKM, SB, SHL, SL, EC, MC, SC and HB performed experiments; ANKM, SB, SHL, SL, AI and SC analyzed data. ANKM wrote the original draft. RNC, AI and SC contributed to manuscript writing, review and editing.

## Supplemental figure legends

**Figure S1: Deficiency mapping using complementation tests with 426.**

Schematic of the region of deficiency lines on chromosome 2. *426* mutants fail to complement deficiency lines indicated by the pink boxes. The gray boxes indicate lines that complement the *426* mutant. Deficiency line *Df(3R)ED6361* for the right arm of chromosome 3 is used here as a negative control.

**Figure S2: Deficiency mapping of *Nckx30c*.**

Schematic of overlapping deficiency lines used for gene mapping. The overlapping region includes the *Nckx30c* gene locus. *426* mutants fail to complement deficiency lines indicated by the pink boxes, implicating cytological region 30C6–30C7 (black dotted line). The phenotype was scored using a yes/no method where a genotype showing a paralytic phenotype was scored as yes, and the one not showing a phenotype was scored as no.

**Figure S3: Deficiency mapping using additional complementation tests with *Nckx30c^426^.*** Schematic of the region of additional deficiency lines on chromosome 2 spanning the region identified as *Nckx30c*. The pink boxes indicate deficiency lines that failed to complement *Nckx30c^426^*.

**Figure S4: cDNA sequencing and alignment of Nckx30c and predicted helix region.**

**(A)** cDNA sequence of *Nckx30c* was acquired from database (Flybase.org) and then it was compared with the cDNA sequences from control *y^1^w^67c23^* and mutant *Nckx30c^426^*flies. The alignment against cDNA from the database shows a change in adenine (A) to cytosine (C) at nucleotide position 1126 in the mutant *Nckx30c^426^*, indicated by the black discontinued line. Subsequent translated amino acid sequence alignment shows a change in threonine (T) to proline (P) at residue position 376. **(B)** Predicted helix region (tool: PSIPRED) shows that the T might be present in the helix region (marked by a green box). Amino acid residues highlighted in yellow represent the beta strand, amino acid residues highlighted in pink represent the alpha helix and amino acid residues highlighted in gray represent irregular coils.

**Figure S5: *Nckx30c* mutants exhibit neurodegeneration.**

Representative 5-μm thin paraffin sections at approximately midbrain of 26-day old control *y^1^w^67c23^* (n=3) and *Nckx30c^426^* (n=3) mutant flies. The presence of holes (arrowheads) is indicative of neurodegeneration. Images were taken at 20x magnification using a light microscope.

**Figure S6: *Nckx30c^426^* mutants exhibit shorter lifespan.**

Lifespan curves of both female and male *Nckx30c^426^*mutants, *y^1^w^67c23^* controls and heterozygous *Nckx30c^426^*/+ flies (*Nckx30c^426^* crossed with *y^1^w^67c23^*) at 25°C. *Nckx30c^426^* flies of both sexes display a significantly shorter lifespan in comparison to *y^1^w^67c23^* and *Nckx30c^426^*/+ flies. Numbers in parentheses indicate sample size. Log-Rank (Mantel-Cox); *P ≤ 0.05, ****P ≤ 0.0001. P values are reported in the **Supplemental datasheet 1.**

**Figure S7: Locomotor behavior analysis of *Nckx30c* compound heterozygotes at 22°C.** Automated open-field behavior tracking to measure distance traveled at 22°C in female and male flies at 5 and 30 days of age, respectively. Numbers on the Y axis indicate the sample size of each compound heterozygote. The horizontal lines across each graph indicate the mean distance traveled for the homozygous mutants *w^1118^*; *Nckx30c^426^, w^1118^*; *Nckx30c^MB07279^, and w^1118^*; *Nckx30c^MB06102^*. A one-way ANOVA (rank-sum post hoc test) was performed to compare each compound heterozygote with *w^1118^*; the symbols over each box represent statistical significance. \**P* ≤ 0.05, **P ≤ 0.01, \*\*\**P* ≤ 0.001, *ns* = not significant. P values are reported in the Supplemental datasheet 1.

**Figure S8: Locomotor behavior analysis of *Nckx30c* compound heterozygotes at 39°C.** Automated open-field behavior tracking to measure distance traveled at 39°C in female and male flies at 5 and 30 days of age, respectively. Numbers on the Y axis indicate the sample size of each compound heterozygote. The horizontal lines across each graph indicate the mean distance traveled for the homozygous mutants *w^1118^*; *Nckx30c^426^, w^1118^*; *Nckx30c^MB07279^, and w^1118^*; *Nckx30c^MB06102^*. A one-way ANOVA (rank-sum post hoc test) was performed to compare each compound heterozygote with *w^1118^*; the symbols over each box represent statistical significance. **P ≤ 0.01, \*\*\**P* ≤ 0.001. P values are reported in the **Supplemental datasheet 1.**

**Figure S9: Linear fit of giant fiber latency as a function of temperature.**

Scatter plot of the log10 of giant fiber latency as a function of temperature. Lines indicate best fits to the equation: log_10_(GF latency) = *A* (ΔT) + *B*. ΔT indicates the difference from initial temperature (20 °C) and the fit parameter *A* and *B* represent the temperature dependence coefficient and latency at initial temperature respectively. For w*^1118^*; *Nckx30c^426^*: *A =*-5.48 x 10^-3^ ± 0.26 x 10^-3^, *B =* 0.29 ± 2.9 x 10^-3^. For *w^1118^*: *A =*-7.84 x 10^-3^ ± 0.28 x 10^-3^, *B =* 0.28 ± 3.3 x 10-3.

**Figure S10: RT-qPCR analysis for the verification of Nckx30c knockdown.**

RT-qPCR analysis of relative expression of *Nckx30c* gene in heads of *elav-Gal4;UAS-dcr2>*+, *elav-Gal4;UAS-dcr2>UAS-Nckx30cRNAi*, *repo-Gal4>*+ and *repo-Gal4>UAS-Nckx30cRNAi* flies. Flies with neuronal *Nckx30c* knockdown (*elav-Gal4;UAS-dcr2>UAS-Nckx30cRNAi*) show lower expression of *Nckx30c* than *elav-Gal4;UAS-dcr2>*+ controls. Flies with glial *Nckx30c* knockdown (*repo-Gal4>UAS-Nckx30cRNAi*) also show lower expression of *Nckx30c* in comparison to *repo-Gal4>*+ controls. Independent experiment number: n=1. Gene expression data were normalized to the *Rp49* gene as an endogenous control, and each knockdown was also normalized to its cell-type control. *elav-Gal4;UAS-dcr2>UAS-Nckx30cRNAi* was normalized to *elav-Gal4;UAS-dcr2>*+ and *repo-Gal4>UAS-Nckx30cRNAi* was normalized to *repo-Gal4>*+.

**Figure S11: RT-qPCR analysis of calcium regulating genes.**

RT-qPCR analysis of relative expression of *MCU and PMCA* genes, respectively, in heads of *w^1118^;Nckx30c^426^*, *w^1118^;Nckx30c^MB07279^ and w^1118^;Nckx30c^MB06102^*flies compared to *w^1118^* controls. All samples were from 7-days old male flies. Independent experiment number: n=6. Mean ± SEM; one-way ANOVA with Dunnett’s post-hoc test; *ns* = not significant. All gene expression data were normalized to *Rp49* gene and shown as relative to *w^1118^*. P values are reported in the **Supplemental datasheet 1.**

**Supplemental Table 1: List of primers used in this study.**

**Supplemental datasheet 1: P-values from statistical analysis in all figures. Supplemental datasheet 2: Raw data for all figures.**

## References

Al-Khannaq, M., & Lytton, J. (2022). Regulation of K(+)-Dependent Na(+)/Ca(2+)-Exchangers (NCKX). Int J Mol Sci, 24(1). 10.3390/ijms24010598

Allen, M. J., Godenschwege, T. A., Tanouye, M. A., & Phelan, P. (2006). Making an escape: development and function of the Drosophila giant fibre system. Semin Cell Dev Biol, 17(1), 31–41. 10.1016/j.semcdb.2005.11.011

Altimimi, H. F., & Schnetkamp, P. P. (2007). Na+/Ca2+-K+ exchangers (NCKX): functional properties and physiological roles. Channels (Austin), 1(2), 62–69. 10.4161/chan.4366

Amatniek, J. C., Hauser, W. A., DelCastillo-Castaneda, C., Jacobs, D. M., Marder, K., Bell, K., Albert, M., Brandt, J., & Stern, Y. (2006). Incidence and Predictors of Seizures in Patients with Alzheimer’s Disease. Epilepsia, 47(5), 867–872. 10.1111/j.1528-1167.2006.00554.x

Arganda-Carreras, I., Fernández-González, R., Muñoz-Barrutia, A., & Ortiz-De-Solorzano, C. (2010). 3D reconstruction of histological sections: Application to mammary gland tissue. Microscopy Research and Technique, 73(11), 1019–1029. 10.1002/jemt.20829

Bagur, R., & Hajnóczky, G. (2017). Intracellular Ca(2+) Sensing: Its Role in Calcium Homeostasis and Signaling. Mol Cell, 66(6), 780–788. 10.1016/j.molcel.2017.05.028

Baines, C. P., Kaiser, R. A., Purcell, N. H., Blair, N. S., Osinska, H., Hambleton, M. A., Brunskill, E. W., Sayen, M. R., Gottlieb, R. A., Dorn, G. W., Robbins, J., & Molkentin, J. D. (2005). Loss of cyclophilin D reveals a critical role for mitochondrial permeability transition in cell death. Nature, 434(7033), 658–662. 10.1038/nature03434

Beramendi, A., Peron, S., Casanova, G., Reggiani, C., & Cantera, R. (2007). Neuromuscular junction in abdominal muscles of Drosophila melanogaster during adulthood and aging. J Comp Neurol, 501(4), 498–508. 10.1002/cne.21253

Bernardi, P., Krauskopf, A., Basso, E., Petronilli, V., Blachly-Dyson, E., Di Lisa, F., & Forte, M. A. (2006). The mitochondrial permeability transition from in vitro artifact to disease target. FEBS J, 273(10), 2077–2099. 10.1111/j.1742-4658.2006.05213.x

Berridge, M. J. (1998). Neuronal Calcium Signaling. Neuron, 21(1), 13–26. 10.1016/S0896-6273(00)80510-3

Bertoni, M., Kiefer, F., Biasini, M., Bordoli, L., & Schwede, T. (2017). Modeling protein quaternary structure of homo-and hetero-oligomers beyond binary interactions by homology. Sci Rep, 7(1), 10480. 10.1038/s41598-017-09654-8

Bienert, S., Waterhouse, A., de Beer, T. A., Tauriello, G., Studer, G., Bordoli, L., & Schwede, T. (2017). The SWISS-MODEL Repository-new features and functionality. Nucleic Acids Res, 45(D1), D313–d319. 10.1093/nar/gkw1132

Bilen, J., & Bonini, N. M. (2005). Drosophila as a model for human neurodegenerative disease. Annu Rev Genet, 39, 153–171. 10.1146/annurev.genet.39.110304.095804

Bkaily, G., & Jacques, D. (2023). Calcium Homeostasis, Transporters, and Blockers in Health and Diseases of the Cardiovascular System. Int J Mol Sci, 24(10). 10.3390/ijms24108803

Brini, M., Calì, T., Ottolini, D., & Carafoli, E. (2014). Neuronal calcium signaling: function and dysfunction. Cell Mol Life Sci, 71(15), 2787–2814. 10.1007/s00018-013-1550-7

Britzolaki, A., Saurine, J., Flaherty, E., Thelen, C., & Pitychoutis, P. M. (2018). The SERCA2: A Gatekeeper of Neuronal Calcium Homeostasis in the Brain. Cell Mol Neurobiol, 38(5), 981–994. 10.1007/s10571-018-0583-8

Budnik, V., Zhong, Y., & Wu, C. F. (1990). Morphological plasticity of motor axons in Drosophila mutants with altered excitability. J Neurosci, 10(11), 3754–3768. 10.1523/jneurosci.10-11-03754.1990

Burg, M. G., & Wu, C. F. (2012). Mechanical and temperature stressor-induced seizure-and-paralysis behaviors in Drosophila bang-sensitive mutants. J Neurogenet, 26(2), 189–197. 10.3109/01677063.2012.690011

Cao, Y., Chtarbanova, S., Petersen, A. J., & Ganetzky, B. (2013). Dnr1 mutations cause neurodegeneration in *Drosophila* by activating the innate immune response in the brain. Proceedings of the National Academy of Sciences, 110(19), E1752–E1760. doi:10.1073/pnas.1306220110

Carraro, M., & Bernardi, P. (2023). The mitochondrial permeability transition pore in Ca(2+) homeostasis. Cell Calcium, 111, 102719. 10.1016/j.ceca.2023.102719

Chaudhuri, D., Artiga, D. J., Abiria, S. A., & Clapham, D. E. (2016). Mitochondrial calcium uniporter regulator 1 (MCUR1) regulates the calcium threshold for the mitochondrial permeability transition. Proc Natl Acad Sci U S A, 113(13), E1872–1880. 10.1073/pnas.1602264113

Chi, W., Iyengar, A. S. R., Albersen, M., Bosma, M., Verhoeven-Duif, N. M., Wu, C.-F., & Zhuang, X. (2019). Pyridox (am) ine 5’-phosphate oxidase deficiency induces seizures in Drosophila melanogaster. Human Molecular Genetics, 28(18), 3126–3136. 10.1093/hmg/ddz143

Choi, S., Quan, X., Bang, S., Yoo, H., Kim, J., Park, J., Park, K. S., & Chung, J. (2017). Mitochondrial calcium uniporter in Drosophila transfers calcium between the endoplasmic reticulum and mitochondria in oxidative stress-induced cell death. J Biol Chem, 292(35), 14473–14485. 10.1074/jbc.M116.765578

Chorna, T., & Hasan, G. (2012). The genetics of calcium signaling in Drosophila melanogaster. Biochimica et Biophysica Acta (BBA) - General Subjects, 1820(8), 1269–1282. 10.1016/j.bbagen.2011.11.002

Collins, C. A., & DiAntonio, A. (2007). Synaptic development: insights from Drosophila. Curr Opin Neurobiol, 17(1), 35–42. 10.1016/j.conb.2007.01.001

Cuomo, O., Sirabella, R., Boscia, F., Casamassa, A., Lytton, J., Annunziato, L., & Pignataro, G. (2022). K(+)-Dependent Na(+)/Ca(2+) Exchanger Isoform 2, Nckx2, Takes Part in the Neuroprotection Elicited by Ischemic Preconditioning in Brain Ischemia. Int J Mol Sci, 23(13). 10.3390/ijms23137128

Dare, S. S., Merlo, E., Rodriguez Curt, J., Ekanem, P. E., Hu, N., & Berni, J. (2020). Drosophila para (bss) Flies as a Screening Model for Traditional Medicine: Anticonvulsant Effects of Annona senegalensis. Front Neurol, 11, 606919. 10.3389/fneur.2020.606919

Dolphin, A. C. (2021). Functions of Presynaptic Voltage-gated Calcium Channels. Function (Oxf), 2(1), zqaa027. 10.1093/function/zqaa027

Elkins, T., Ganetzky, B., & Wu, C.-F. (1986). A Drosophila Mutation that Eliminates a Calcium-Dependent Potassium Current. Proceedings of the National Academy of Sciences of the United States of America, 83(21), 8415–8419. http://www.jstor.org/stable/28597

Engel, J. E., & Wu, C.-F. (1998). Genetic Dissection of Functional Contributions of Specific Potassium Channel Subunits in Habituation of an Escape Circuit in <em>Drosophila</em>. The Journal of Neuroscience, 18(6), 2254–2267. 10.1523/jneurosci.18-06-02254.1998

Engel, J. E., & Wu, C. F. (1996). Altered habituation of an identified escape circuit in Drosophila memory mutants. J Neurosci, 16(10), 3486–3499. 10.1523/jneurosci.16-10-03486.1996

Erhardt, B., Marcora, M. S., Frenkel, L., Bochicchio, P. A., Bodin, D. H., Silva, B. A., Farías, M. I., Allo, M., Höcht, C., Ferrari, C. C., Pitossi, F. J., & Leal, M. C. (2021). Plasma membrane calcium ATPase downregulation in dopaminergic neurons alters cellular physiology and motor behaviour in Drosophila melanogaster. Eur J Neurosci, 54(6), 5915–5931. 10.1111/ejn.15401

Feng, B., & Chen, Z. (2016). Generation of febrile seizures and subsequent epileptogenesis. Neuroscience bulletin, 32(5), 481–492.

Florentinus-Mefailoski, A., Bowden, P., Scheltens, P., Killestein, J., Teunissen, C., & Marshall, J. G. (2021). The plasma peptides of Alzheimer’s disease. Clin Proteomics, 18(1), 17. 10.1186/s12014-021-09320-2

Folk, D. G., Hoekstra, L. A., & Gilchrist, G. W. (2007). Critical thermal maxima in knockdown-selected Drosophila: are thermal endpoints correlated? Journal of Experimental Biology, 210(15), 2649–2656. 10.1242/jeb.003350

Gadhave, D. G., Sugandhi, V. V., Jha, S. K., Nangare, S. N., Gupta, G., Singh, S. K., Dua, K., Cho, H., Hansbro, P. M., & Paudel, K. R. (2024). Neurodegenerative disorders: Mechanisms of degeneration and therapeutic approaches with their clinical relevance. Ageing Res Rev, 99, 102357. 10.1016/j.arr.2024.102357

Ganetzky, B., & Wu, C. F. (1986). Neurogenetics of membrane excitability in Drosophila. Annu Rev Genet, 20, 13–44. 10.1146/annurev.ge.20.120186.000305

Gao, H. M., & Hong, J. S. (2008). Why neurodegenerative diseases are progressive: uncontrolled inflammation drives disease progression. Trends Immunol, 29(8), 357–365. 10.1016/j.it.2008.05.002

Gevedon, O., Bolus, H., Lye, S. H., Schmitz, K., Fuentes-González, J., Hatchell, K., Bley, L., Pienaar, J., Loewen, C., & Chtarbanova, S. (2019). In Vivo Forward Genetic Screen to Identify Novel Neuroprotective Genes in Drosophila melanogaster. J Vis Exp(149). 10.3791/59720

Gleichmann, M., & Mattson, M. P. (2011). Neuronal calcium homeostasis and dysregulation. Antioxid Redox Signal, 14(7), 1261–1273. 10.1089/ars.2010.3386

Gourmaud, S., Stewart, D. A., Irwin, D. J., Roberts, N., Barbour, A. J., Eberwine, G., O’Brien, W. T., Vassar, R., Talos, D. M., & Jensen, F. E. (2022). The role of mTORC1 activation in seizure-induced exacerbation of Alzheimer’s disease. Brain, 145(1), 324–339. 10.1093/brain/awab268

Gramates, L. S., & Budnik, V. (1999). Assembly and maturation of the Drosophila larval neuromuscular junction. Int Rev Neurobiol, 43, 93–117. 10.1016/s0074-7742(08)60542-5

Guex, N., & Peitsch, M. C. (1997). SWISS-MODEL and the Swiss-Pdb Viewer: An environment for comparative protein modeling. ELECTROPHORESIS, 18(15), 2714–2723. 10.1002/elps.1150181505

Haug-Collet, K., Pearson, B., Webel, R., Szerencsei, R. T., Winkfein, R. J., Schnetkamp, P. P., & Colley, N. J. (1999). Cloning and characterization of a potassium-dependent sodium/calcium exchanger in Drosophila. J Cell Biol, 147(3), 659–670. 10.1083/jcb.147.3.659

Horigane, S.-i., Ozawa, Y., Yamada, H., & Takemoto-Kimura, S. (2019). Calcium signalling: a key regulator of neuronal migration. The Journal of Biochemistry, 165(5), 401–409. 10.1093/jb/mvz012

Iyengar, A., Imoehl, J., Ueda, A., Nirschl, J., & Wu, C. F. (2012). Automated quantification of locomotion, social interaction, and mate preference in Drosophila mutants. J Neurogenet, 26(3-4), 306–316. 10.3109/01677063.2012.729626

Iyengar, A., Ruan, H., & Wu, C. F. (2022). Distinct Aging-Vulnerable and-Resilient Trajectories of Specific Motor Circuit Functions in Oxidation-and Temperature-Stressed Drosophila. eNeuro, 9(1). 10.1523/eneuro.0443-21.2021

Iyengar, A., & Wu, C. F. (2014). Flight and seizure motor patterns in Drosophila mutants: simultaneous acoustic and electrophysiological recordings of wing beats and flight muscle activity. J Neurogenet, 28(3-4), 316–328. 10.3109/01677063.2014.957827

Iyengar, A., & Wu, C. F. (2021). Fly seizure EEG: field potential activity in the Drosophila brain. J Neurogenet, 35(3), 295–305. 10.1080/01677063.2021.1950714

Jalloul, A. H., Szerencsei, R. T., Rogasevskaia, T. P., & Schnetkamp, P. P. M. (2018). SLC24A Family (K+-Dependent Na+-Ca2+ Exchanger, NCKX). In S. Choi (Ed.), Encyclopedia of Signaling Molecules (pp. 4994–5002). Springer International Publishing. 10.1007/978-3-319-67199-4_101860

Jalloul, A. H., Szerencsei, R. T., Rogasevskaia, T. P., & Schnetkamp, P. P. M. (2020). Structure-function relationships of K(+)-dependent Na(+)/Ca(2+) exchangers (NCKX). Cell Calcium, 86, 102153. 10.1016/j.ceca.2019.102153

Juel, V. C. (2012). Evaluation of neuromuscular junction disorders in the electromyography laboratory. Neurol Clin, 30(2), 621–639. 10.1016/j.ncl.2011.12.012

Kang, K., Bauer, P. J., Kinjo, T. G., Szerencsei, R. T., Bönigk, W., Winkfein, R. J., & Schnetkamp, P. P. M. (2003). Assembly of Retinal Rod or Cone Na+/Ca2+-K+Exchanger Oligomers with cGMP-Gated Channel Subunits as Probed with Heterologously Expressed cDNAs. Biochemistry, 42(15), 4593–4600. 10.1021/bi027276z

Kim, E. Y., Kang, K. H., & Koh, H. (2018). Cyclophilin 1 (Cyp1) mutation ameliorates oxidative stress-induced defects in a Drosophila DJ-1 null mutant. Biochem Biophys Res Commun, 505(3), 823–829. 10.1016/j.bbrc.2018.10.014

Klein, P., Carrazana, E., Glauser, T., Herman, B. P., Penovich, P., Rabinowicz, A. L., & Sutula, T. P. (2025). Do Seizures Damage the Brain?-Cumulative Effects of Seizures and Epilepsy: A 2025 Perspective. Epilepsy Curr, 15357597251331927. 10.1177/15357597251331927

Krebs, J. (2022). Structure, Function and Regulation of the Plasma Membrane Calcium Pump in Health and Disease. Int J Mol Sci, 23(3). 10.3390/ijms23031027

Kroll, J. R., Wong, K. G., Siddiqui, F. M., & Tanouye, M. A. (2015). Disruption of Endocytosis with the Dynamin Mutant shibirets1 Suppresses Seizures in Drosophila. Genetics, 201(3), 1087–1102. 10.1534/genetics.115.177600

Landaverde, S., Sleep, M., Lacoste, A., Tan, S., Schuback, R., Reiter, L. T., & Iyengar, A. (2024). Glial expression of Drosophila UBE3A causes spontaneous seizures that can be modulated by 5-HT signaling. Neurobiology of Disease, 200, 106651. 10.1016/j.nbd.2024.106651

Lee, J., Iyengar, A., & Wu, C. F. (2019). Distinctions among electroconvulsion-and proconvulsant-induced seizure discharges and native motor patterns during flight and grooming: quantitative spike pattern analysis in flight muscles. Journal of Neurogenetics, 33(2), 125–142. 10.1080/01677063.2019.1581188

Lee, J., & Wu, C. F. (2002). Electroconvulsive seizure behavior in Drosophila: analysis of the physiological repertoire underlying a stereotyped action pattern in bang-sensitive mutants. J Neurosci, 22(24), 11065–11079. 10.1523/jneurosci.22-24-11065.2002

Lee, J., & Wu, C. F. (2010). Orchestration of stepwise synaptic growth by K+ and Ca2+ channels in Drosophila. J Neurosci, 30(47), 15821–15833. 10.1523/jneurosci.3448-10.2010

Lehmann, L., Lo, A., Knox, K. M., & Barker-Haliski, M. (2021). Alzheimer’s Disease and Epilepsy: A Perspective on the Opportunities for Overlapping Therapeutic Innovation. Neurochem Res, 46(8), 1895–1912. 10.1007/s11064-021-03332-y

Loewen, C., Boekhoff-Falk, G., Ganetzky, B., & Chtarbanova, S. (2018). A Novel Mutation in Brain Tumor Causes Both Neural Over-Proliferation and Neurodegeneration in Adult Drosophila. G3 Genes|Genomes|Genetics, 8(10), 3331–3346. 10.1534/g3.118.200627

Mahapatra, C., Thakkar, R., & Kumar, R. (2024). Modulatory Impact of Oxidative Stress on Action Potentials in Pathophysiological States: A Comprehensive Review. Antioxidants (Basel), 13(10). 10.3390/antiox13101172

Majlish, A. N. K., Bourgeois, S., Lye, S. H., Landaverde, S., Cytron, E., Cline, M., Bolus, H., Correll, R. N., Iyengar, A., & Chtarbanova, S. (2025). Supplemental material for Majlish ANK et al.10.17605/OSF.IO/YQDVM

McGuffin, L. J., Bryson, K., & Jones, D. T. (2000). The PSIPRED protein structure prediction server. Bioinformatics, 16(4), 404–405. 10.1093/bioinformatics/16.4.404

Melom, J. E., & Littleton, J. T. (2013). Mutation of a NCKX Eliminates Glial Microdomain Calcium Oscillations and Enhances Seizure Susceptibility. The Journal of Neuroscience, 33(3), 1169–1178. 10.1523/jneurosci.3920-12.2013

Menon, K. P., Carrillo, R. A., & Zinn, K. (2013). Development and plasticity of the Drosophila larval neuromuscular junction. Wiley Interdiscip Rev Dev Biol, 2(5), 647–670. 10.1002/wdev.108

Mozolewski, P., Jeziorek, M., Schuster, C. M., Bading, H., Frost, B., & Dobrowolski, R. (2021). The role of nuclear Ca2+ in maintaining neuronal homeostasis and brain health. Journal of Cell Science, 134(8). 10.1242/jcs.254904

Negi, D., Granak, S., Shorter, S., O’Leary, V. B., Rektor, I., & Ovsepian, S. V. (2023). Molecular Biomarkers of Neuronal Injury in Epilepsy Shared with Neurodegenerative Diseases. Neurotherapeutics, 20(3), 767–778. 10.1007/s13311-023-01355-7

Nichols, E., Steinmetz, J. D., Vollset, S. E., Fukutaki, K., Chalek, J., Abd-Allah, F., Abdoli, A., Abualhasan, A., Abu-Gharbieh, E., Akram, T. T., Al Hamad, H., Alahdab, F., Alanezi, F. M., Alipour, V., Almustanyir, S., Amu, H., Ansari, I., Arabloo, J., Ashraf, T., Astell-Burt, T., Ayano, G., Ayuso-Mateos, J. L., Baig, A. A., Barnett, A., Barrow, A., Baune, B. T., Béjot, Y., Bezabhe, W. M. M., Bezabih, Y. M., Bhagavathula, A. S., Bhaskar, S., Bhattacharyya, K., Bijani, A., Biswas, A., Bolla, S. R., Boloor, A., Brayne, C., Brenner, H., Burkart, K., Burns, R. A., Cámera, L. A., Cao, C., Carvalho, F., Castro-de-Araujo, L. F. S., Catalá-López, F., Cerin, E., Chavan, P. P., Cherbuin, N., Chu, D.-T., Costa, V. M., Couto, R. A. S., Dadras, O., Dai, X., Dandona, L., Dandona, R., De la Cruz-Góngora, V., Dhamnetiya, D., Dias da Silva, D., Diaz, D., Douiri, A., Edvardsson, D., Ekholuenetale, M., El Sayed, I., El-Jaafary, S. I., Eskandari, K., Eskandarieh, S., Esmaeilnejad, S., Fares, J., Faro, A., Farooque, U., Feigin, V. L., Feng, X., Fereshtehnejad, S.-M., Fernandes, E., Ferrara, P., Filip, I., Fillit, H., Fischer, F., Gaidhane, S., Galluzzo, L., Ghashghaee, A., Ghith, N., Gialluisi, A., Gilani, S. A., Glavan, I.-R., Gnedovskaya, E. V., Golechha, M., Gupta, R., Gupta, V. B., Gupta, V. K., Haider, M. R., Hall, B. J., Hamidi, S., Hanif, A., Hankey, G. J., Haque, S., Hartono, R. K., Hasaballah, A. I., Hasan, M. T., Hassan, A., Hay, S. I., Hayat, K., Hegazy, M. I., Heidari, G., Heidari-Soureshjani, R., Herteliu, C., Househ, M., Hussain, R., Hwang, B.-F., Iacoviello, L., Iavicoli, I., Ilesanmi, O. S., Ilic, I. M., Ilic, M. D., Irvani, S. S. N., Iso, H., Iwagami, M., Jabbarinejad, R., Jacob, L., Jain, V., Jayapal, S. K., Jayawardena, R., Jha, R. P., Jonas, J. B., Joseph, N., Kalani, R., Kandel, A., Kandel, H., Karch, A., Kasa, A. S., Kassie, G. M., Keshavarz, P., Khan, M. A. B., Khatib, M. N., Khoja, T. A. M., Khubchandani, J., Kim, M. S., Kim, Y. J., Kisa, A., Kisa, S., Kivimäki, M., Koroshetz, W. J., Koyanagi, A., Kumar, G. A., Kumar, M., Lak, H. M., Leonardi, M., Li, B., Lim, S. S., Liu, X., Liu, Y., Logroscino, G., Lorkowski, S., Lucchetti, G., Lutzky Saute, R., Magnani, F. G., Malik, A. A., Massano, J., Mehndiratta, M. M., Menezes, R. G., Meretoja, A., Mohajer, B., Mohamed Ibrahim, N., Mohammad, Y., Mohammed, A., Mokdad, A. H., Mondello, S., Moni, M. A. A., Moniruzzaman, M., Mossie, T. B., Nagel, G., Naveed, M., Nayak, V. C., Neupane Kandel, S., Nguyen, T. H., Oancea, B., Otstavnov, N., Otstavnov, S. S., Owolabi, M. O., Panda-Jonas, S., Pashazadeh Kan, F., Pasovic, M., Patel, U. K., Pathak, M., Peres, M. F. P., Perianayagam, A., Peterson, C. B., Phillips, M. R., Pinheiro, M., Piradov, M. A., Pond, C. D., Potashman, M. H., Pottoo, F. H., Prada, S. I., Radfar, A., Raggi, A., Rahim, F., Rahman, M., Ram, P., Ranasinghe, P., Rawaf, D. L., Rawaf, S., Rezaei, N., Rezapour, A., Robinson, S. R., Romoli, M., Roshandel, G., Sahathevan, R., Sahebkar, A., Sahraian, M. A., Sathian, B., Sattin, D., Sawhney, M., Saylan, M., Schiavolin, S., Seylani, A., Sha, F., Shaikh, M. A., Shaji, K. S., Shannawaz, M., Shetty, J. K., Shigematsu, M., Shin, J. I., Shiri, R., Silva, D. A. S., Silva, J. P., Silva, R., Singh, J. A., Skryabin, V. Y., Skryabina, A. A., Smith, A. E., Soshnikov, S., Spurlock, E. E., Stein, D. J., Sun, J., Tabarés-Seisdedos, R., Thakur, B., Timalsina, B., Tovani-Palone, M. R., Tran, B. X., Tsegaye, G. W., Valadan Tahbaz, S., Valdez, P. R., Venketasubramanian, N., Vlassov, V., Vu, G. T., Vu, L. G., Wang, Y.-P., Wimo, A., Winkler, A. S., Yadav, L., Yahyazadeh Jabbari, S. H., Yamagishi, K., Yang, L., Yano, Y., Yonemoto, N., Yu, C., Yunusa, I., Zadey, S., Zastrozhin, M. S., Zastrozhina, A., Zhang, Z.-J., Murray, C. J. L., & Vos, T. (2022). Estimation of the global prevalence of dementia in 2019 and forecasted prevalence in 2050: an analysis for the Global Burden of Disease Study 2019. The Lancet Public Health, 7(2), e105–e125. 10.1016/S2468-2667(21)00249-8

Nikoletopoulou, V., & Tavernarakis, N. (2012). Calcium homeostasis in aging neurons [Review]. Frontiers in Genetics, Volume 3 - 2012. 10.3389/fgene.2012.00200

Noebels, J. (2011). A perfect storm: Converging paths of epilepsy and Alzheimer’s dementia intersect in the hippocampal formation. Epilepsia, 52 Suppl 1(Suppl 1), 39–46. 10.1111/j.1528-1167.2010.02909.x

Okonechnikov, K., Golosova, O., Fursov, M., & team, t. U. (2012). Unipro UGENE: a unified bioinformatics toolkit. Bioinformatics, 28(8), 1166–1167. 10.1093/bioinformatics/bts091

Palladino, M. J., Hadley, T. J., & Ganetzky, B. (2002). Temperature-sensitive paralytic mutants are enriched for those causing neurodegeneration in Drosophila. Genetics, 161(3), 1197–1208. 10.1093/genetics/161.3.1197

Park, J.-H., Park, I., Youm, E. M., Lee, S., Park, J.-H., Lee, J., Lee, D. Y., Byun, M. S., Lee, J. H., Yi, D., Chung, S. J., Park, K. W., Choi, N., Kim, S. Y., Yoon, W., An, H., Kim, K. w., Choi, S. H., Jeong, J. H., Kim, E.-J., Kang, H., Lee, J., Kim, Y., Lee, E. A., Seo, S. W., Na, D. L., & Kim, J.-W. (2021). Novel Alzheimer’s disease risk variants identified based on whole-genome sequencing of APOE ε4 carriers. Translational Psychiatry, 11(1), 296. 10.1038/s41398-021-01412-9

Pavlidis, P., & Tanouye, M. A. (1995). Seizures and failures in the giant fiber pathway of Drosophila bang-sensitive paralytic mutants. J Neurosci, 15(8), 5810–5819. 10.1523/jneurosci.15-08-05810.1995

Pchitskaya, E., Popugaeva, E., & Bezprozvanny, I. (2018). Calcium signaling and molecular mechanisms underlying neurodegenerative diseases. Cell Calcium, 70, 87–94. 10.1016/j.ceca.2017.06.008

Peña-Bautista, C., Casas-Fernández, E., Vento, M., Baquero, M., & Cháfer-Pericás, C. (2020). Stress and neurodegeneration. Clin Chim Acta, 503, 163–168. 10.1016/j.cca.2020.01.019

Pizzo, P., Drago, I., Filadi, R., & Pozzan, T. (2012). Mitochondrial Ca2+ homeostasis: mechanism, role, and tissue specificities. Pflügers Archiv - European Journal of Physiology, 464(1), 3–17. 10.1007/s00424-012-1122-y

Pratt, J., De Vito, G., Narici, M., & Boreham, C. (2021). Neuromuscular Junction Aging: A Role for Biomarkers and Exercise. J Gerontol A Biol Sci Med Sci, 76(4), 576–585. 10.1093/gerona/glaa207

Purushotham, M., Tashrifwala, F., Jena, R., Vudugula, S. A., Patil, R. S., & Agrawal, A. (2022). The Association Between Alzheimer’s Disease and Epilepsy: A Narrative Review. Cureus, 14(10), e30195. 10.7759/cureus.30195

Rossetto, M. G., Zanarella, E., Orso, G., Scorzeto, M., Megighian, A., Kumar, V., Delgado-Escueta, A. V., & Daga, A. (2011). Defhc1.1, a homologue of the juvenile myoclonic gene EFHC1, modulates architecture and basal activity of the neuromuscular junction in Drosophila. Hum Mol Genet, 20(21), 4248–4257. 10.1093/hmg/ddr352

Ruan, H., Ueda, A., Xing, X., Wan, X., Strub, B., Mukai, S., Certel, K., Green, D., Belozerov, K., Yao, W. D., Johnson, W., Jung-Ching Lin, J., Hilliker, A. J., & Wu, C. F. (2017). Generation and characterization of new alleles of quiver (qvr) that encodes an extracellular modulator of the Shaker potassium channel. J Neurogenet, 31(4), 325–336. 10.1080/01677063.2017.1393076

Schnetkamp, P. P. (2013). The SLC24 gene family of Na⁺/Ca²⁺-K⁺ exchangers: from sight and smell to memory consolidation and skin pigmentation. Mol Aspects Med, 34(2-3), 455–464. 10.1016/j.mam.2012.07.008

Scholtens, L. H., Pijnenburg, R., de Lange, S. C., Huitinga, I., & van den Heuvel, M. P. (2022). Common Microscale and Macroscale Principles of Connectivity in the Human Brain. J Neurosci, 42(20), 4147–4163. 10.1523/jneurosci.1572-21.2022

Schwarzer, A., Kim, T. S. Y., Hagen, V., Molday, R. S., & Bauer, P. J. (1997). The Na/Ca-K Exchanger of Rod Photoreceptor Exists as Dimer in the Plasma Membrane. Biochemistry, 36(44), 13667–13676. 10.1021/bi9710232

Sidisky, J. M., & Babcock, D. T. (2020). Visualizing Synaptic Degeneration in Adult Drosophila in Association with Neurodegeneration. J Vis Exp(159). 10.3791/61363

Sidisky, J. M., Weaver, D., Hussain, S., Okumus, M., Caratenuto, R., & Babcock, D. (2021). Mayday sustains trans-synaptic BMP signaling required for synaptic maintenance with age. Elife, 10. 10.7554/eLife.54932

Stafstrom, C. E., & Carmant, L. (2015). Seizures and epilepsy: an overview for neuroscientists. Cold Spring Harb Perspect Med, 5(6). 10.1101/cshperspect.a022426

Stolwijk, J. A., Zhang, X., Gueguinou, M., Zhang, W., Matrougui, K., Renken, C., & Trebak, M. (2016). Calcium Signaling Is Dispensable for Receptor Regulation of Endothelial Barrier Function. J Biol Chem, 291(44), 22894–22912. 10.1074/jbc.M116.756114

Studer, G., Rempfer, C., Waterhouse, A. M., Gumienny, R., Haas, J., & Schwede, T. (2020). QMEANDisCo-distance constraints applied on model quality estimation. Bioinformatics, 36(6), 1765–1771. 10.1093/bioinformatics/btz828

Sun, D., Amiri, M., Meng, Q., Unnithan, R. R., & French, C. (2024). Calcium Signalling in Neurological Disorders, with Insights from Miniature Fluorescence Microscopy. Cells, 14(1). 10.3390/cells14010004

Suzuki, D. T., Grigliatti, T., & Williamson, R. (1971). Temperature-sensitive mutations in Drosophila melanogaster. VII. A mutation (para-ts) causing reversible adult paralysis. Proc Natl Acad Sci U S A, 68(5), 890–893. 10.1073/pnas.68.5.890

Tombini, M., Assenza, G., Ricci, L., Lanzone, J., Boscarino, M., Vico, C., Magliozzi, A., & Di Lazzaro, V. (2021). Temporal Lobe Epilepsy and Alzheimer’s Disease: From Preclinical to Clinical Evidence of a Strong Association. J Alzheimers Dis Rep, 5(1), 243–261. 10.3233/adr-200286

Verbeek, N. E., Wassenaar, M., van Campen, J. S., Sonsma, A., Gunning, B., Knoers, N., Lindhout, D., Jansen, F. E., Leijten, F., Brilstra, E. H., & Kasteleijn-Nolst Trenité, D. (2015). Seizure precipitants in Dravet syndrome: What events and activities are specifically provocative compared with other epilepsies? Epilepsy & Behavior, 47, 39–44. 10.1016/j.yebeh.2015.05.008

von Stockum, S., Basso, E., Petronilli, V., Sabatelli, P., Forte, M. A., & Bernardi, P. (2011). Properties of Ca(2+) transport in mitochondria of Drosophila melanogaster. J Biol Chem, 286(48), 41163–41170. 10.1074/jbc.M111.268375

Vonhoff, F., & Keshishian, H. (2017). In Vivo Calcium Signaling during Synaptic Refinement at the Drosophila Neuromuscular Junction. J Neurosci, 37(22), 5511–5526. 10.1523/jneurosci.2922-16.2017

Wang, M., Zhang, H., Liang, J., Huang, J., Wu, T., & Chen, N. (2025). Calcium signaling hypothesis: A non-negligible pathogenesis in Alzheimer’s disease. J Adv Res. 10.1016/j.jare.2025.01.007

Wang, P., Saraswati, S., Guan, Z., Watkins, C. J., Wurtman, R. J., & Littleton, J. T. (2004). A Drosophila temperature-sensitive seizure mutant in phosphoglycerate kinase disrupts ATP generation and alters synaptic function. J Neurosci, 24(19), 4518–4529. 10.1523/jneurosci.0542-04.2004

Wang, Y., Moussian, B., Schaeffeler, E., Schwab, M., & Nies, A. T. (2018). The fruit fly Drosophila melanogaster as an innovative preclinical ADME model for solute carrier membrane transporters, with consequences for pharmacology and drug therapy. Drug Discovery Today, 23(10), 1746–1760. 10.1016/j.drudis.2018.06.002

Waterhouse, A., Bertoni, M., Bienert, S., Studer, G., Tauriello, G., Gumienny, R., Heer, F. T., de Beer, T. A. P., Rempfer, C., Bordoli, L., Lepore, R., & Schwede, T. (2018). SWISS-MODEL: homology modelling of protein structures and complexes. Nucleic Acids Res, 46(W1), W296–W303. 10.1093/nar/gky427

Webel, R., Haug-Collet, K., Pearson, B., Szerencsei, R. T., Winkfein, R. J., Schnetkamp, P. P., & Colley, N. J. (2002). Potassium-dependent sodium-calcium exchange through the eye of the fly. Ann N Y Acad Sci, 976, 300–314. 10.1111/j.1749-6632.2002.tb04753.x

Williamson, R., Kaplan, W. D., & Dagan, D. (1974). A fly’s leap from paralysis. Nature, 252(5480), 224–226. 10.1038/252224a0

Winkfein, R. J., Pearson, B., Ward, R., Szerencsei, R. T., Colley, N. J., & Schnetkamp, P. P. (2004). Molecular characterization, functional expression and tissue distribution of a second NCKX Na+/Ca2+-K+ exchanger from Drosophila. Cell Calcium, 36(2), 147–155. 10.1016/j.ceca.2004.01.021

Xie, W., Koppula, S., Kale, M. B., Ali, L. S., Wankhede, N. L., Umare, M. D., Upaganlawar, A. B., Abdeen, A., Ebrahim, E. E., El-Sherbiny, M., Behl, T., Shen, B., & Singla, R. K. (2024). Unraveling the nexus of age, epilepsy, and mitochondria: exploring the dynamics of cellular energy and excitability. Front Pharmacol, 15, 1469053. 10.3389/fphar.2024.1469053

Young, K., & Morrison, H. (2018). Quantifying Microglia Morphology from Photomicrographs of Immunohistochemistry Prepared Tissue Using ImageJ. J Vis Exp(136). 10.3791/57648

Zhang, Y., Sharma, S., & Lytton, J. (2015). Anatomical evidence for a non-synaptic influence of the K+-dependent Na+/Ca2+-exchanger, NCKX2, on hippocampal plasticity. Neuroscience, 310, 372–388. 10.1016/j.neuroscience.2015.09.049

Zhou, X., Chen, Z., Xiao, L., Zhong, Y., Liu, Y., Wu, J., & Tao, H. (2022). Intracellular calcium homeostasis and its dysregulation underlying epileptic seizures. Seizure, 103, 126–136. 10.1016/j.seizure.2022.11.007

