## Supplementary figures and images for "*Nckx30c*, a *Drosophila* K^+^-dependent Na^+^/Ca^2+^ exchanger, regulates temperature-sensitive convulsions and age-related neurodegeneration"

### Supplemental Figure S1

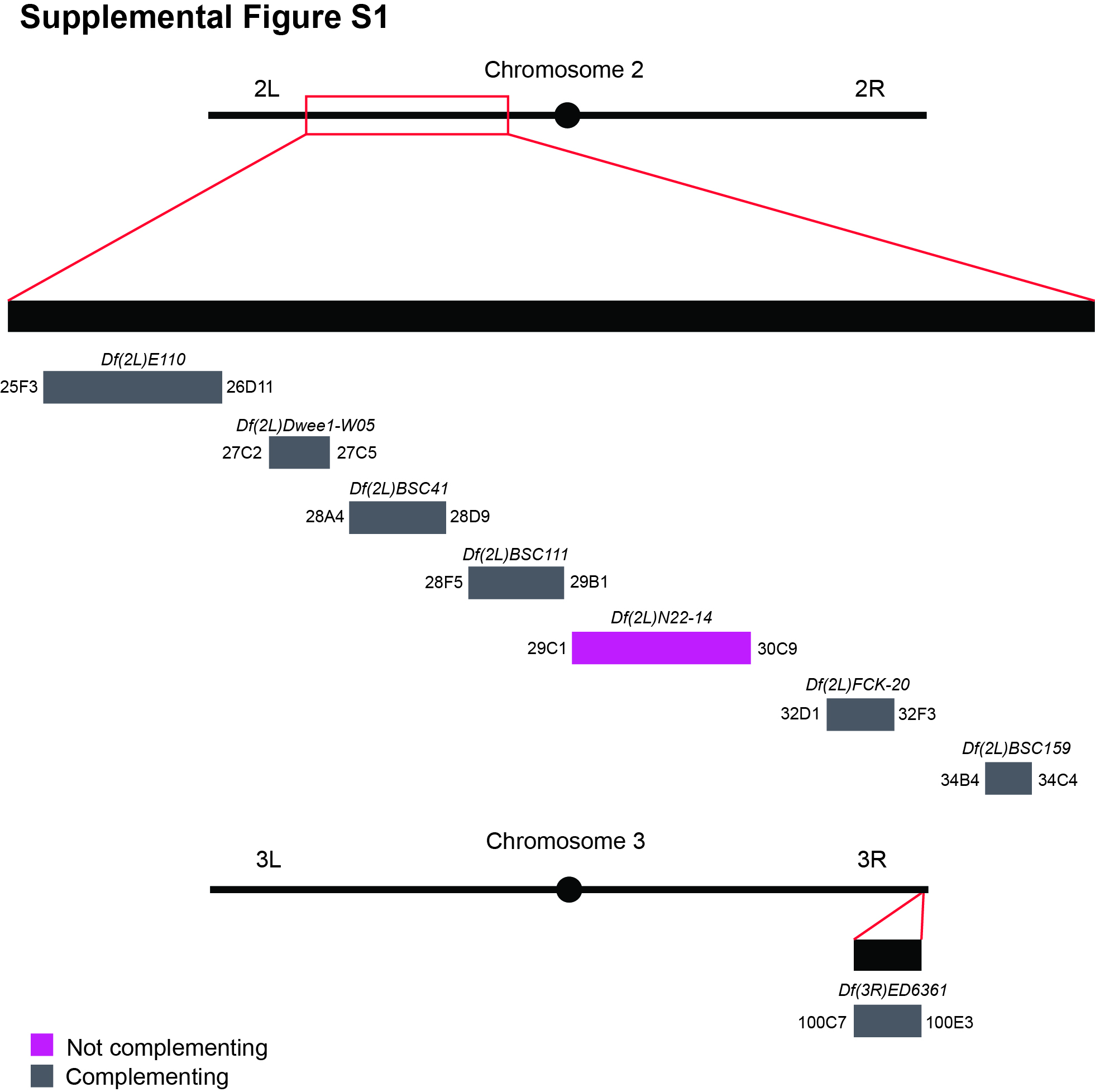

### Supplemental Figure S2

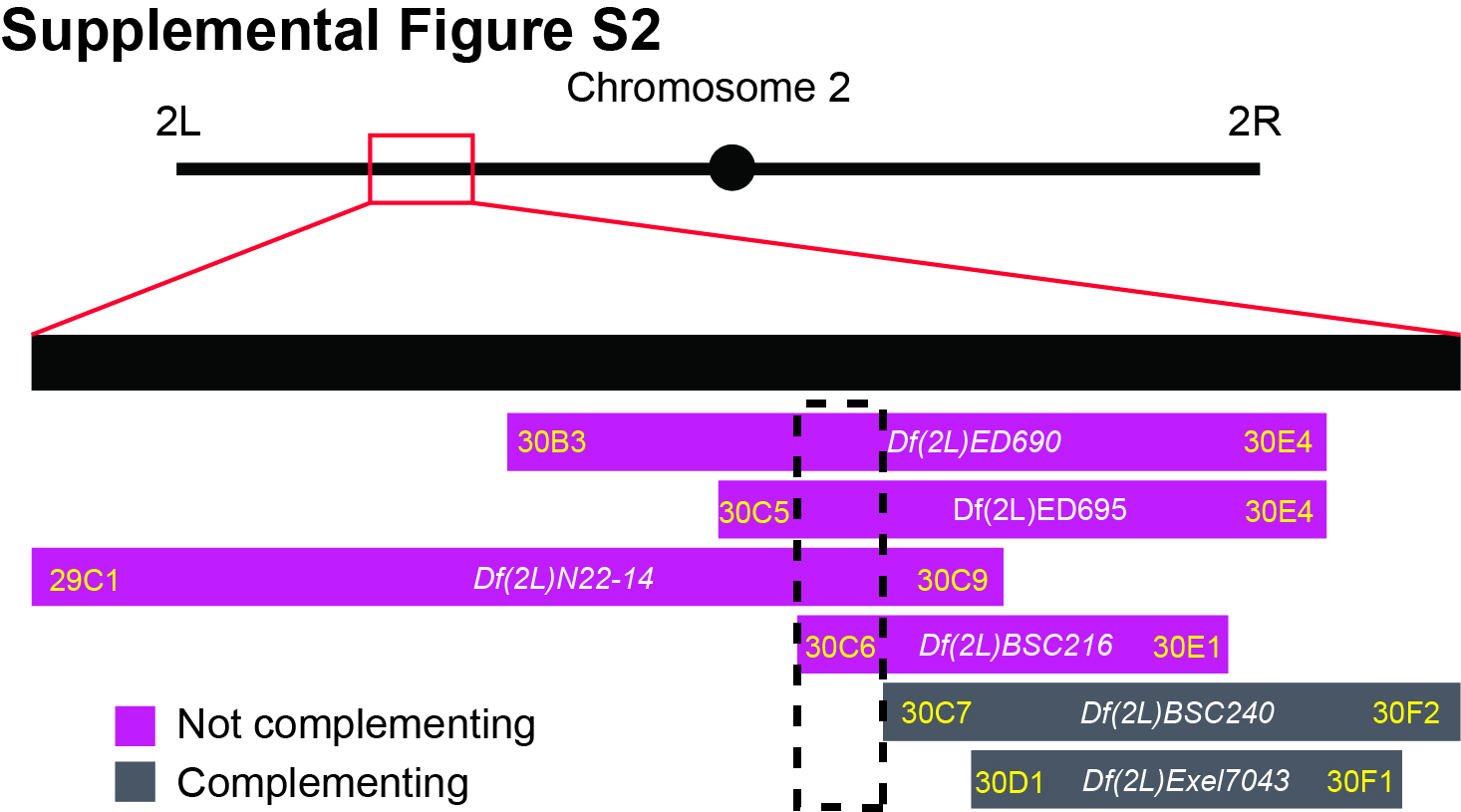

### Supplemental Figure S3

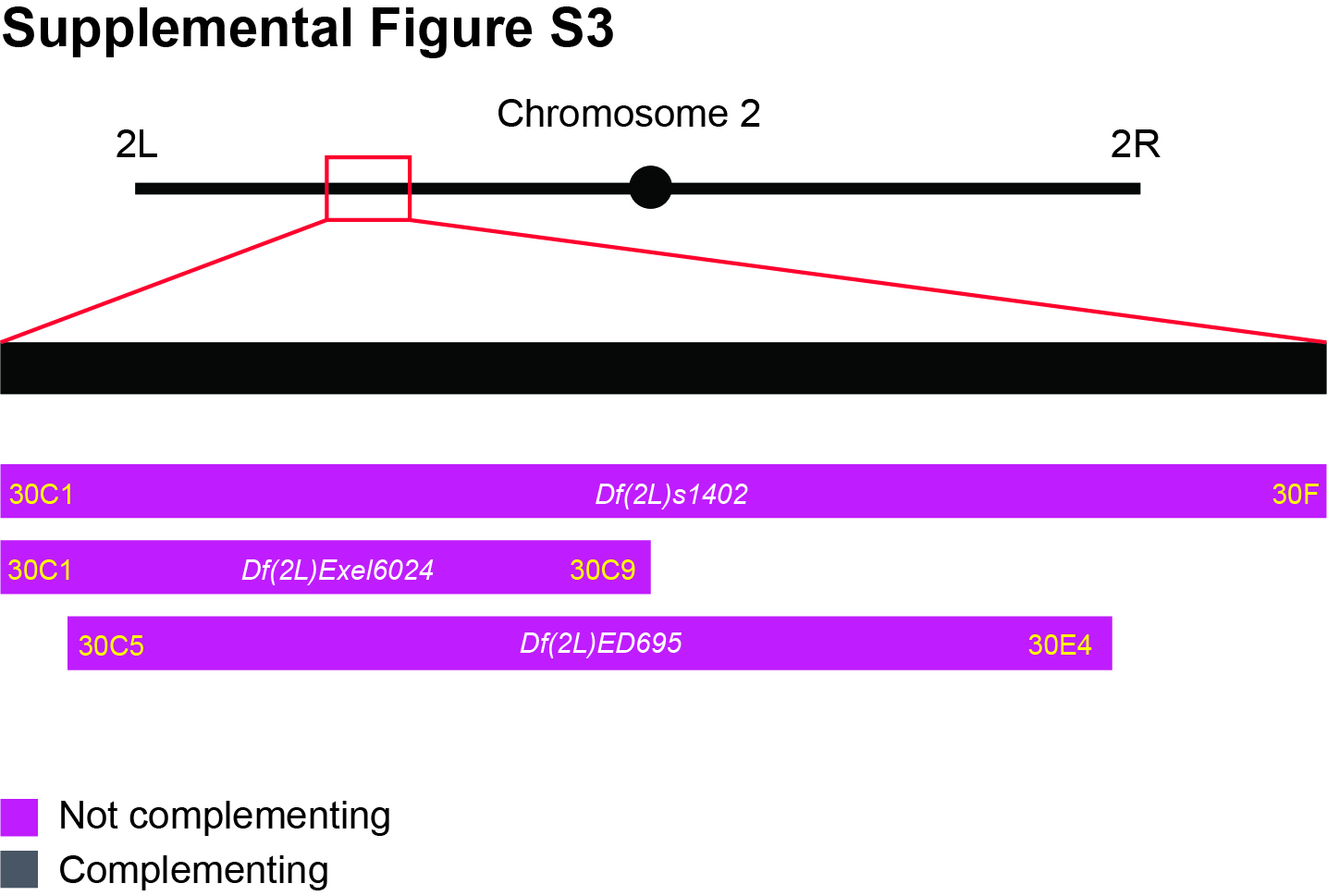

### Supplemental Figure S4

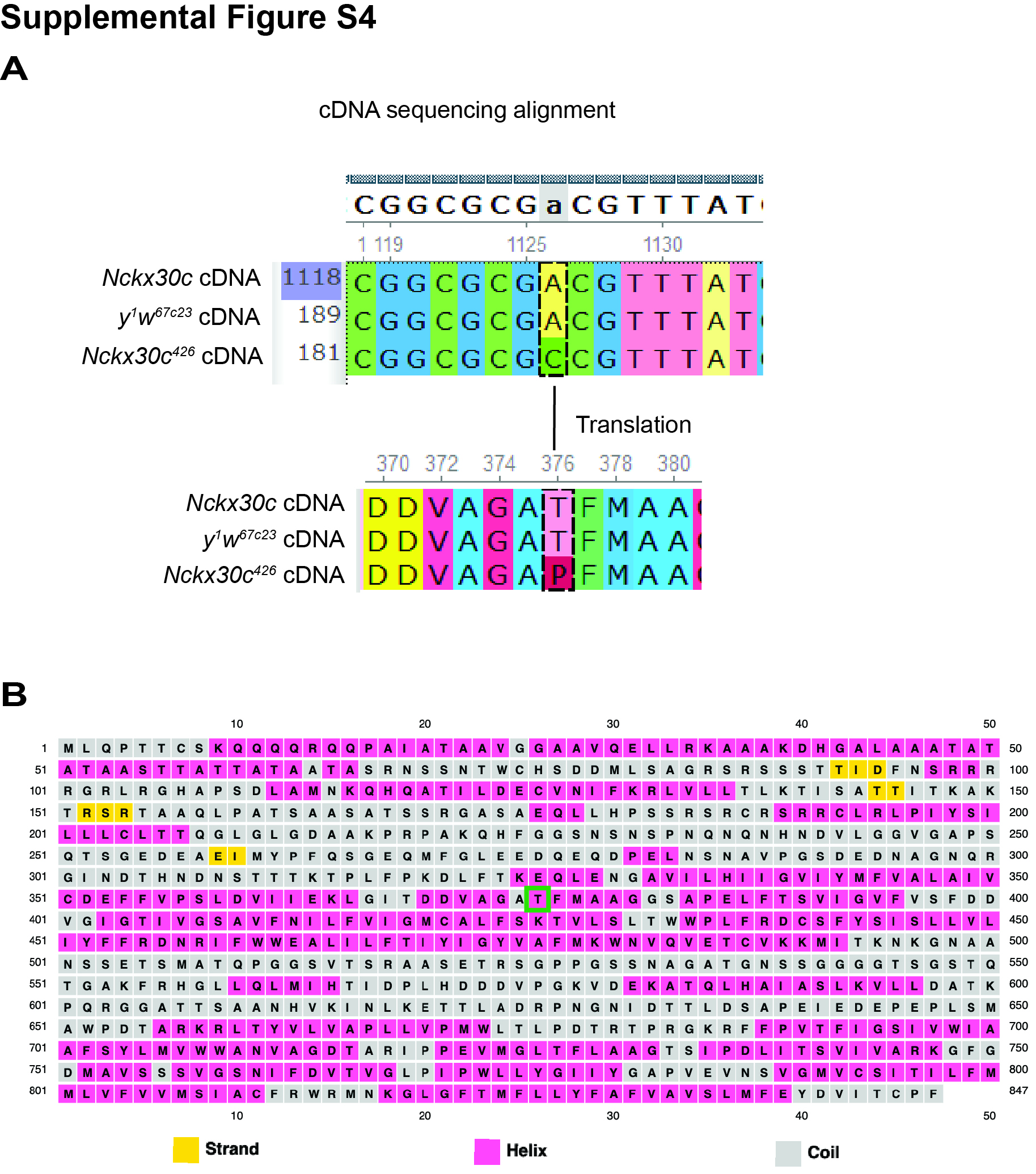

### Supplemental Figure S5

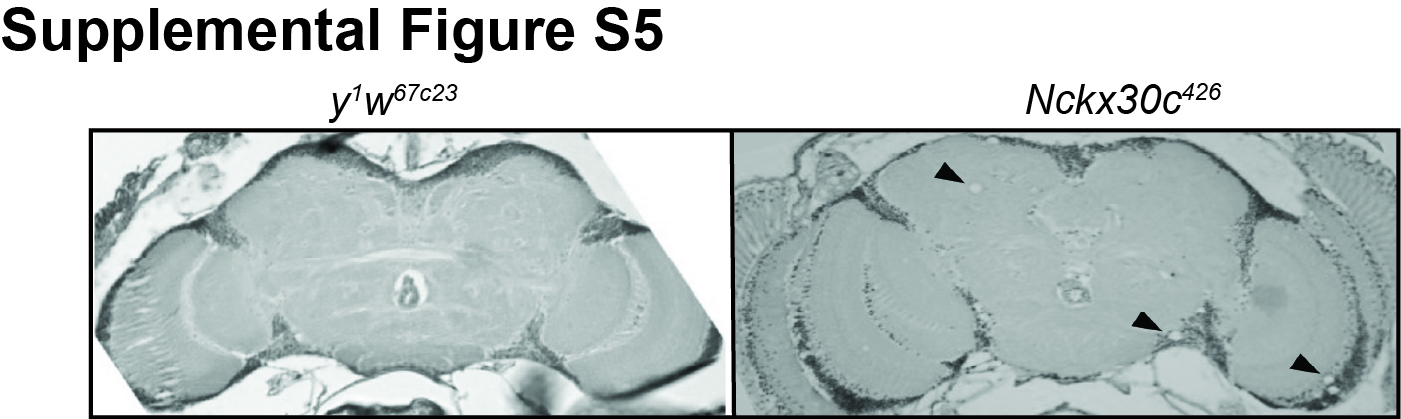

### Supplemental Figure S6

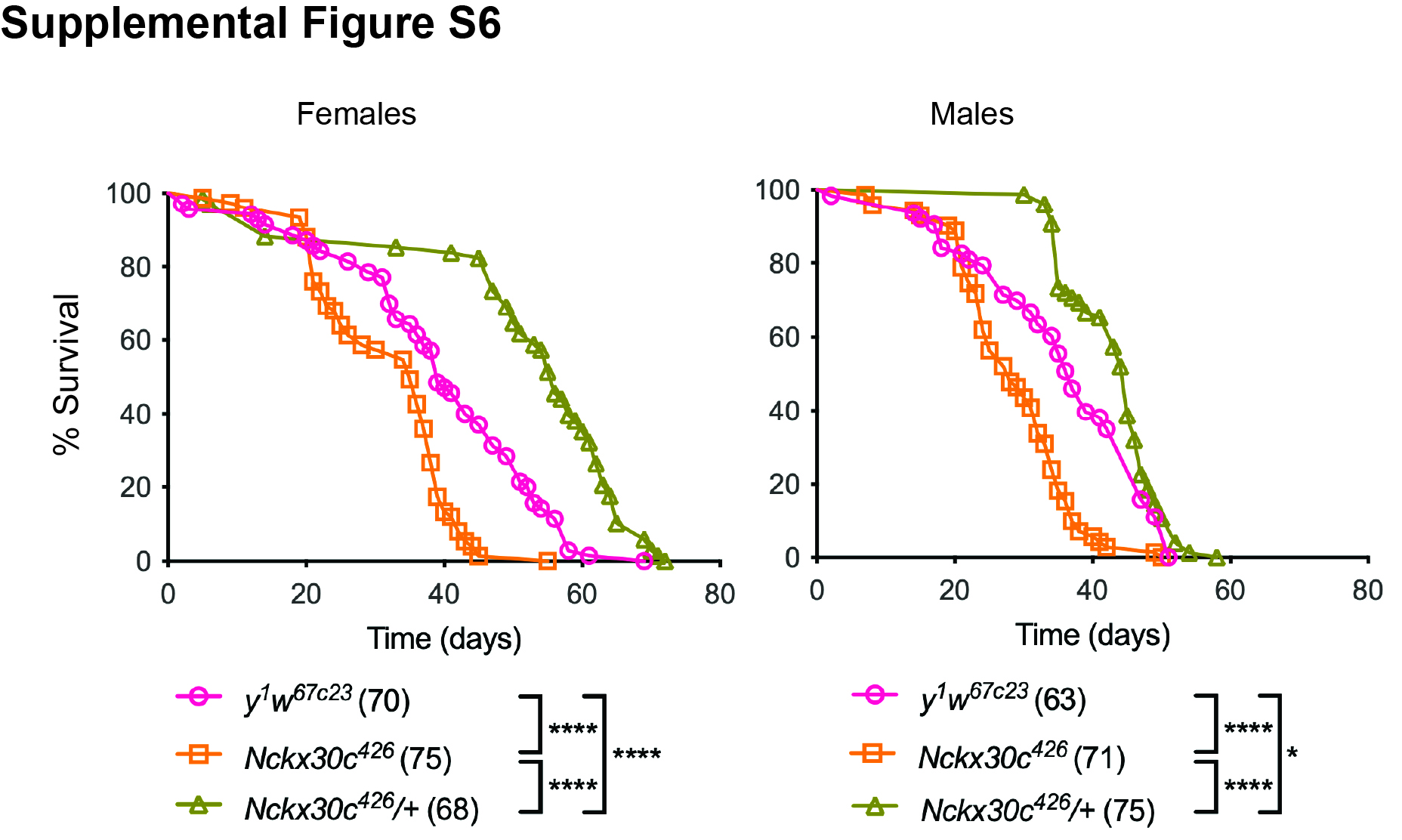

### Supplemental Figure S7

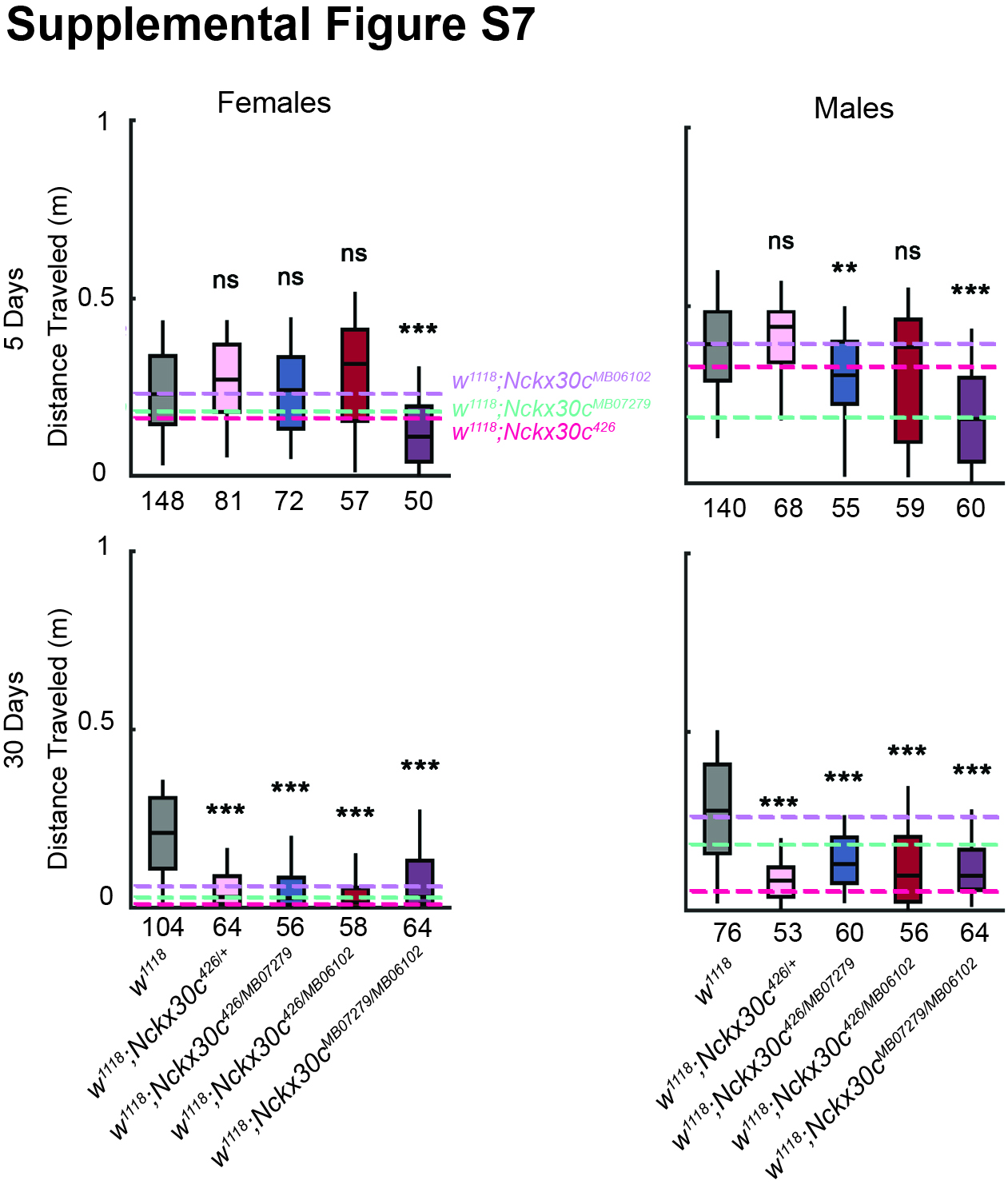

### Supplemental Figure S8

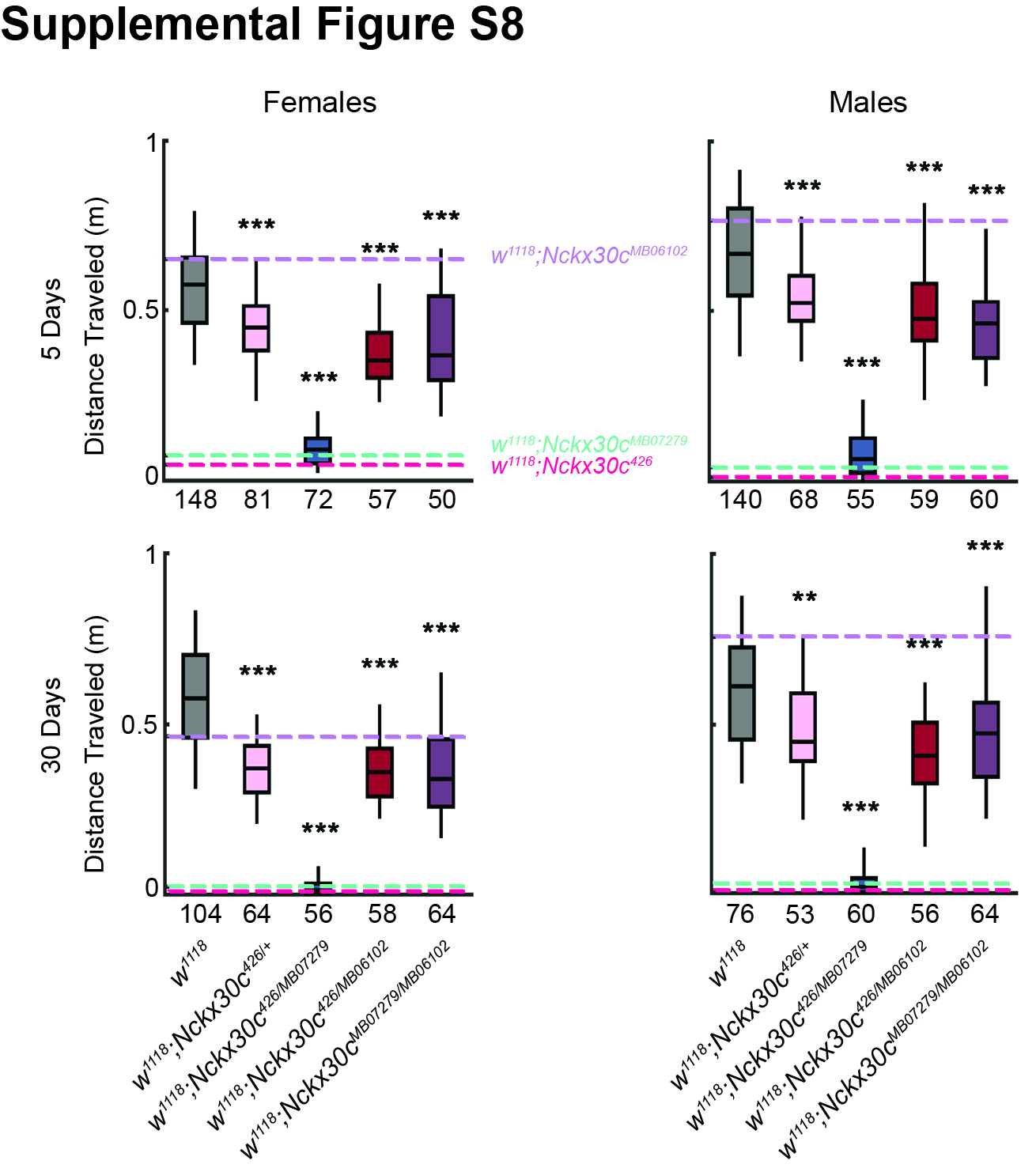

### Supplemental Figure S9

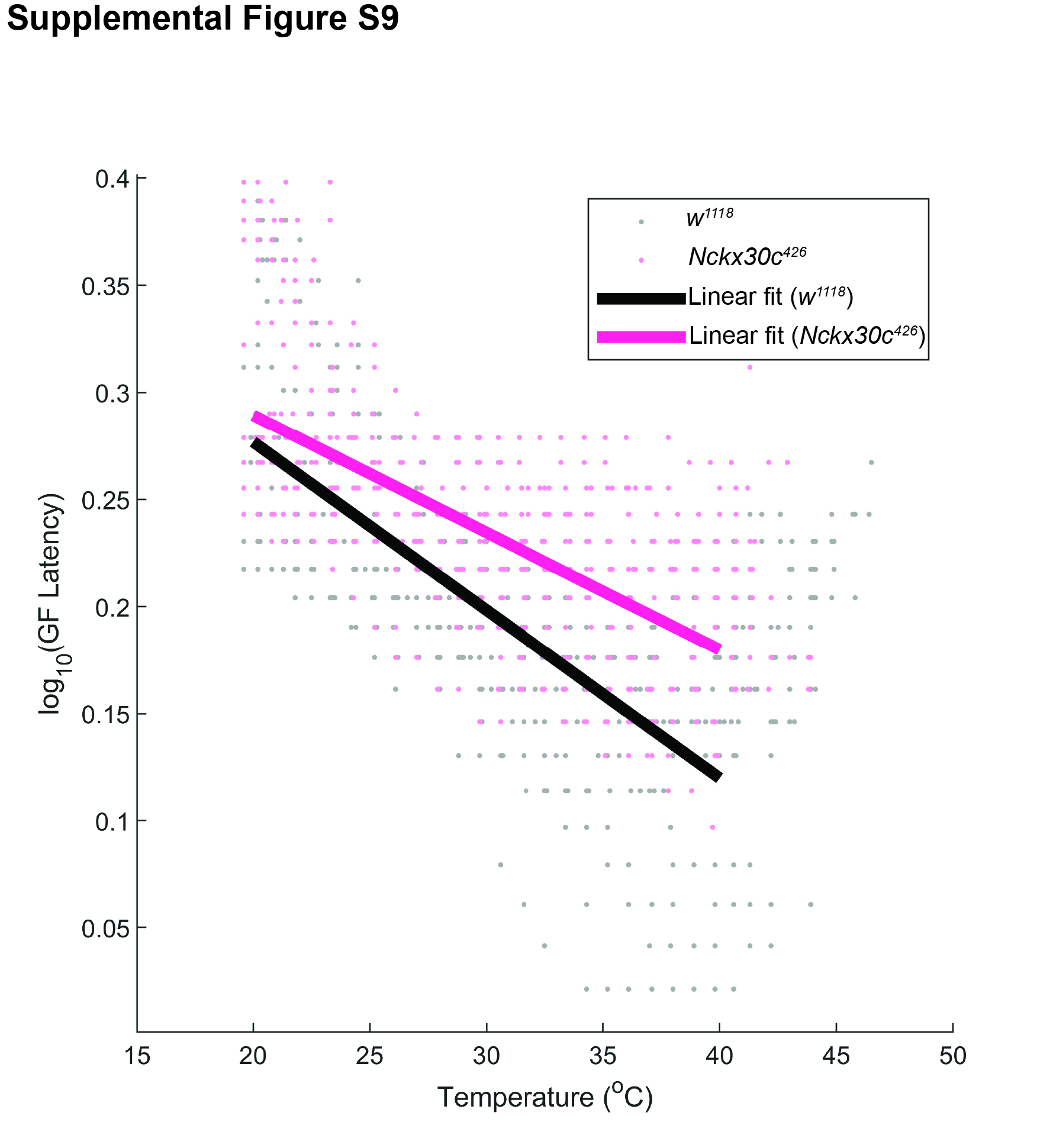

### Supplemental Figure S10

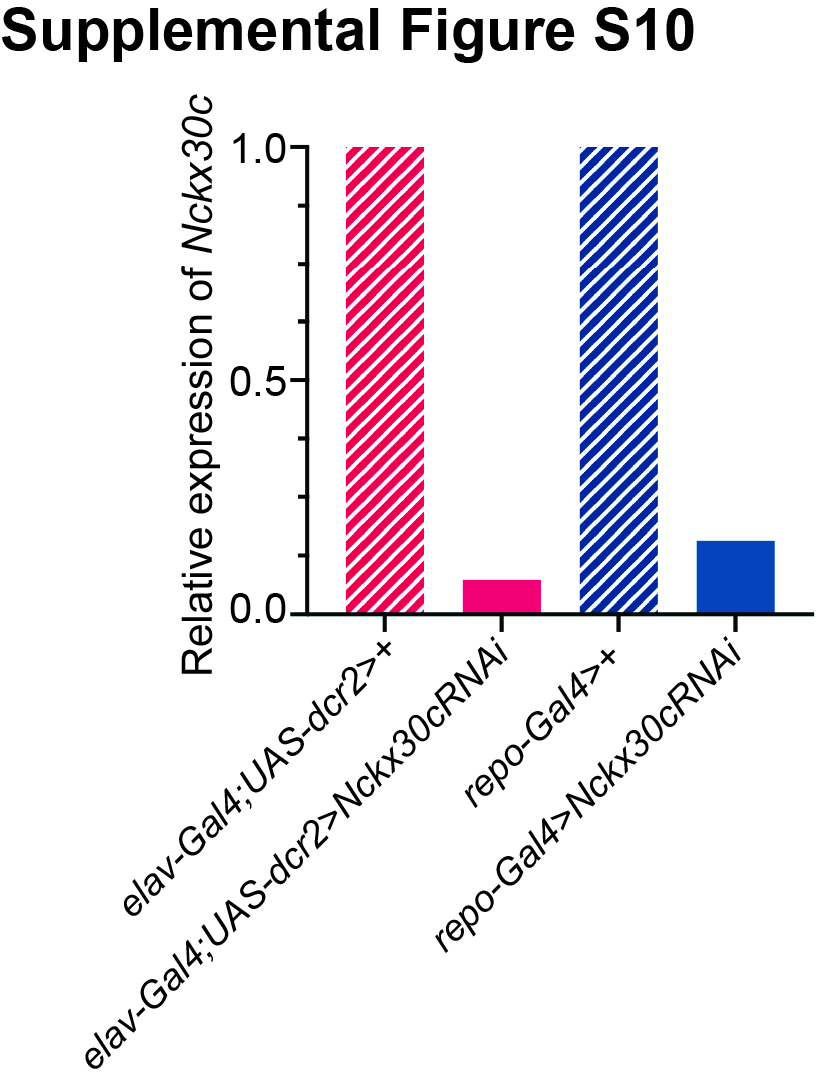

### Supplemental Figure S11

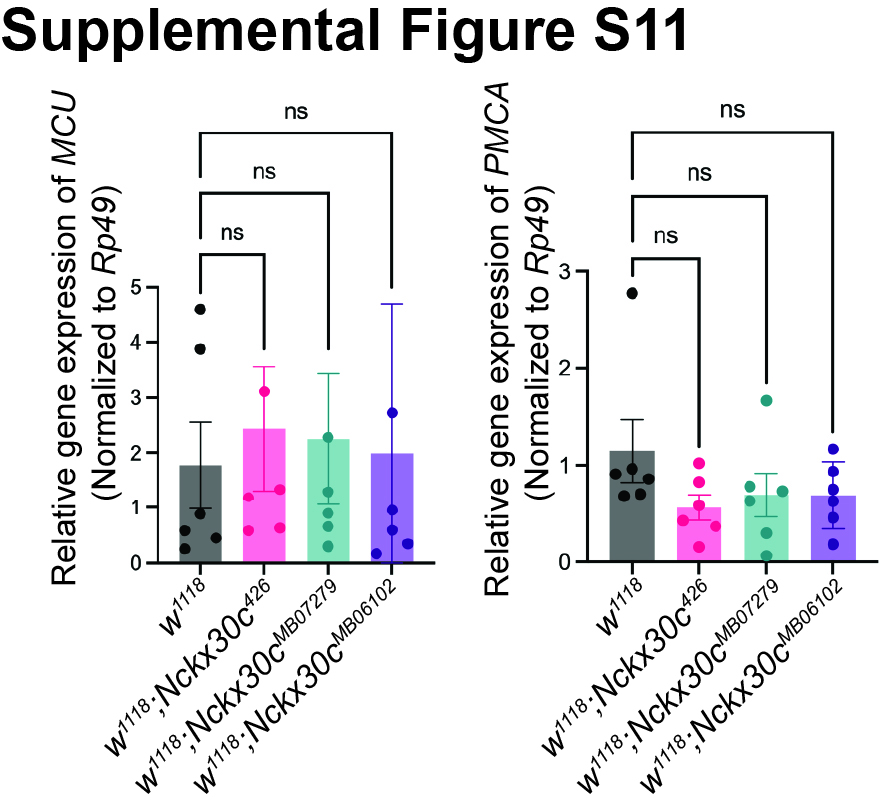
