## Supplemental Table 1 for "*Nckx30c*, a *Drosophila* K^+^-dependent Na^+^/Ca^2+^ exchanger, regulates temperature-sensitive convulsions and age-related neurodegeneration"

**Supplemental Table 1: List of primers used in this study**

| **Target name** | **Primer sequence** |
| --- | --- |
| Exon 1  (PCR) | FW: 5’ – *TGGCGTGAAAAATACCACAAGA* – *3’*  RV: 5’ – CACGACGAACTGATTACCGT – 3’ |
| Exon 2  (PCR) | FW: 5’ – *GCGATGGAAAATTCCTCGAAAA* – *3’*  RV: 5’ –CTCTTTCGCTTTAACTTTCGGT – 3’ |
| Exon 3  (PCR) | FW: 5’ – *CCGATAAACCCGATACCGAAAC* – *3’*  RV: 5’ –*GGTCTCGAAGGGTTGGTGTTA* – 3’ |
| Exon 4  (PCR) | FW: 5’ – *CAACACCACCAACACCTCAC* – *3’*  RV: 5’ –*GCTTGCCTAGACTTACAGCA* – 3’ |
| Exon 5  (PCR) | FW: 5’ – *TCTTTCCCGATCCGTATGTGT* – *3’*  RV: 5’ –*CTTACGGGTCTTACAGTCCAA* – 3’ |
| Exon 6  (PCR) | FW: 5’ – *AGTTCGGTCCAAGTCGCTAA* – 3’  RV: 5’ –*CGTCTGGTTTAAGTAAGGGTCT* – 3’ |
| Exon 7-8  (PCR) | FW: 5’ – *ATTCCGCCCGAGGTACAATC* – 3’  RV: 5’ –*TCGGAGTGTGGGTTAATCGA* – 3’ |
| *Nckx30c* cDNA pair 1 | FW: 5’ – ATTGCTCCACCCAAGTAGCC – 3’  RV: 5’ – GCCCACATCATCGAACGACACG – 3’ |
| *Nckx30c* cDNA pair 2 | FW: 5’ – GCATTGGTACGATCGTTGGC – 3’  RV: 5’ – GGTGGTGTCGATGTTCCCAT – 3’ |
| *Nckx30c* cDNA pair 3 | FW: 5’ – TAGTACTCAGACCGGTGCCA – 3’  RV: 5’ – GTGCCACAATCACCGAGGTA – 3’ |
| *Rp49*  (RT-qPCR) | FW: 5’ – AAGAAGCGCACCAAGCACTTCATC– 3’  FW: 5’ – TCTGTTGTCGATACCCTTGGGCTT– 3’ |
| *Nckx30c*  (RT-qPCR) | FW: 5’ – ATGTTGCAGCCAACAACATG – 3’  RV: 5’ – TTAAAAGGGGCACGTGATCA – 3’ |
| PMCA  (RT-qPCR) | FW: 5’ – GGTCGTGAGGGTGTAATGAAG– 3’  RV: 5’ – GTCAGCTTTCGATCCACTTAGG– 3’ |
| *MCU*  (RT-qPCR) | FW: 5’ – ATGAGCTTCAGCAGCACAGA– 3’  RV: 5’ – CTTGAACTGTGACCTCCCGT– 3’ |
